# Galectin-9 interacts with Vamp-3 to regulate cytokine secretion in dendritic cells

**DOI:** 10.1101/2022.05.13.491792

**Authors:** Rui Santalla Méndez, Andrea Rodgers Furones, René Classens, Manon Haverdil, Marta Canela Capdevila, Anne van Duffelen, Cornelia G. Spruijt, Michiel Vermeulen, Martin ter Beest, Annemiek B. van Spriel, Laia Querol Cano

## Abstract

Intracellular vesicle transport is essential for cellular homeostasis and is partially mediated by SNARE proteins. Endosomal trafficking to the plasma membrane ensures cytokine secretion in dendritic cells (DCs) and the initiation of immune responses. Despite its critical importance, the specific molecular agents that regulate DC cytokine secretion are poorly characterised. Galectin-9, a ß-galactoside-binding protein, has emerged as a novel cellular modulator although its exact intracellular roles in regulating (immune) cell homeostasis and vesicle transport are virtually unknown. We investigated galectin-9 function in primary human DCs and report that galectin-9 is essential for intracellular cytokine trafficking to the cell surface. Galectin-9-depleted DCs accumulate cytokine-containing vesicles in the Golgi complex that eventually undergo lysosomal degradation. We observed galectin-9 to molecularly interact with Vamp-3 using immunoprecipitation-mass-spectrometry and identified galectin-9 was required for rerouting Vamp-3-containing endosomes upon DC activation as the underlying mechanism. Overall, this study identifies galectin-9 as a necessary mechanistic component for intracellular trafficking. This may impact our general understanding of vesicle transport and shed new light into the multiple roles galectins play in governing cell function.

## Introduction

Vesicle transport and secretion is essential to maintain intracellular organelle homeostasis and extracellular cell-cell communication. This process is regulated by dynamic changes in the cytoskeleton as well as by fusion proteins such as the soluble N-ethylmaleimide-sensitive factor attachment protein receptor (SNARE) protein superfamily, that, amongst others, includes syntaxins and vesicle-associated membrane proteins (VAMPs) (Hehnly & Stamnes, 2007; Jahn & Scheller, 2006). In conjunction with the cytoskeleton, SNAREs drive vesicle sorting, docking and membrane fusion from the trans-Golgi network (TGN) to the plasma membrane *via* the endosomal machinery (Jahn & Scheller, 2006). The tight regulation of vesicle trafficking and fusion events allows the functioning of complex biological processes such as the secretion of growth factors, neurotransmitters or hormones. In immune cells, SNARE proteins mediate the release of cytokines, chemokines or the delivery of surface components (Revelo et al, 2019). Despite its crucial importance, the molecular mechanisms that regulate SNARE function and govern endosomal trafficking are not fully described.

Dendritic cells (DCs) are unique immune cells specialised in detecting potentially hazardous elements and mounting early stages of innate and adaptive immunity (Banchereau & Steinman, 1998). To function, DCs rely heavily on the timely secretion of specific cytokines that ensure the activation of (immune) cells and the initiation of downstream effector mechanisms (Joffre et al, 2009; Kapsenberg, 2003). DC activation induces context-dependent cytokine transcription (Kapsenberg, 2003; Mazzoni & Segal, 2004). Newly synthesised cytokines are then transported from the endoplasmic reticulum onto the Golgi complex and the vesicle network to the plasma membrane (Murray & Stow, 2014; Stanley & Lacy, 2010; Verboogen et al, 2018). DCs and monocytes use the recycling endosome pathway to sort, traffic and release key cytokines such as tumour necrosis factor (TNF)-α, interleukin (IL)-6, IL-12 and IL-10 (Chiaruttini et al, 2016; Manderson et al, 2007; Murray et al, 2005; Stanley et al, 2012). Several publications have identified Vamp-3, a SNARE protein located in the early and recycling endosomal compartments (Bajno et al, 2000), as a key player for the secretion of these cytokines in DCs through the recycling endosomes pathway (Manderson et al, 2007; Murray et al, 2005; Stanley et al, 2012). Although cytokine secretion has been widely studied, detailed knowledge about the precise molecular components regulating vesicle movement and spatial segregation remains elusive.

Galectins, a family of β-galactoside-binding lectins, have emerged as modulators of cancer, inflammation and autoimmunity (Gordon-Alonso et al, 2018; Liu & Rabinovich, 2010; Vasta et al, 2017). Galectins are located both intracellularly in the cysotol, where they establish carbohydrate-independent protein-protein interactions and extracellularly, where they bind oligosaccharide residues present on membrane glycoproteins, glycolipids and the extracellular matrix creating galectin-mediated lattices (Nabi et al, 2015; Thiemann & Baum, 2016). This confers them the ability to play numerous functions either intracellular or extracellularly, in turn regulating (immune) cell homeostasis (Johannes et al, 2018; Liu et al, 2002). Galectin-9, a ubiquitously expressed tandem-repeat type of galectin, has gained increasing importance due to its multiple roles in governing various cell functions (John & Mishra, 2016). For instance, intracellular galectin-9 has been associated with autophagy in hepatitis B infected cells and to activate AMPK upon lysosomal damage (Jia et al, 2019; Miyakawa et al, 2022). Similarly, galectin-9 has been recently implicated in protein and membrane recycling in phagosome formation in macrophages (Klose et al, 2019). Secreted galectin-9 has been described to play a role in epithelial cell morphology and polarisation *via* its interaction with the Forssman glycosphingolipid (Mishra et al, 2010). Galectin-9 is mostly known for its role in cancer progression as it is reported to participate in tumour cell adhesion, migration and aggregation (Vladoiu et al, 2014). In contrast, virtually nothing is known regarding the involvement of any member of the galectin family in cytokine intracellular trafficking or secretion. The few reports available describe the association of galectin-9 in cytokine secretion in mast cells and macrophages although the underlying molecular mechanisms are not resolved (Kojima et al, 2014; Lv et al, 2017) and galectin-9 function in DC cytokine secretion has not been investigated. To address this, we depleted galectin-9 in human primary DCs and studied its effects on cytokine release. Our experiments revealed that cytokine secretion was significantly impaired in the absence of galectin-9 independently of transcription. We identified a novel molecular interaction between Vamp-3 and galectin-9, which governs the trafficking of Vamp-3 containing vesicles to the plasma membrane upon DC activation as the underlying mechanism. Overall, our work demonstrates that galectin-9 acts as a novel regulator of the cytokine trafficking machinery through controlling Vamp-3 function and subcellular localisation.

## Results

### Galectin-9 is required for cytokine secretion

The transcription and secretion of a large number of cytokines that act as key modulators of innate and adaptive immune responses is induced by immune cell activation. We investigated whether galectin-9 was involved in cytokine secretion using human monocyte derived DCs (moDCs) as model system for immune cell function. moDCs lacking galectin-9 were generated by electroporating them with either specific *galectin-9* (*LGALS9*) siRNA or <u>n</u>on-<u>t</u>argeting (NT) siRNA control (hereby referred to as gal-9 KD and WT cells, respectively) prior to examine their cytokine response to various stimuli. Galectin-9 depletion, both surface-bound and intracellularly, was confirmed by flow cytometry (Supplementary Figure S1A and B) and Western Blot (Supplementary Figure S1C), showing a downregulation of the protein level of approximately 90 %. The involvement of galectin-9 in cytokine secretion was studied in four key cytokines in DCs, namely pro-inflammatory cytokines <u>i</u>nter<u>l</u>eukin (IL)-6 (IL-6) and <u>t</u>umour <u>n</u>ecrosis <u>f</u>actor α (TNFα), the T cell activator cytokine IL-12 and the anti-inflammatory cytokine IL-10. We first determined the optimal secretion time point and stimulus for each of them (Supplementary Figure S2), and observed that IL-6 and TNFα were secreted after stimulation with the bacterial membrane component lipopolysaccharide (LPS) and TLR4 agonist; IL-10 in response to zymosan stimulation (a fungal cell wall extract) and IL-12 in response to R848 (a TLR7 and TLR8 agonist) (Figure S2A). The secretion peak for all cytokines was shown to occur 16 h after treatment, while IL-10 secretion spiked at 24 h instead (Figure S2B). These conditions were then used to measure cytokine secretion in WT and in gal-9 KD moDCs across various independent donors. Depletion of galectin-9 significantly impaired the secretion of all cytokines analysed when compared to WT control cells (Figure 1A). To determine whether galectin-9 depletion resulted in a general protein synthesis and/or trafficking defect upon DC activation, we first studied the expression of plasma membrane proteins known to be enhanced in response to LPS treatment. Flow cytometry performed on activated WT and gal-9 KD cells showed these proteins to be equally upregulated upon incubation with LPS (Figure 1B and 1C). This data indicates that galectin-9 acts specifically in modulating cytokine secretion and does not affect general cellular trafficking to the plasma membrane.

**Figure 1.**
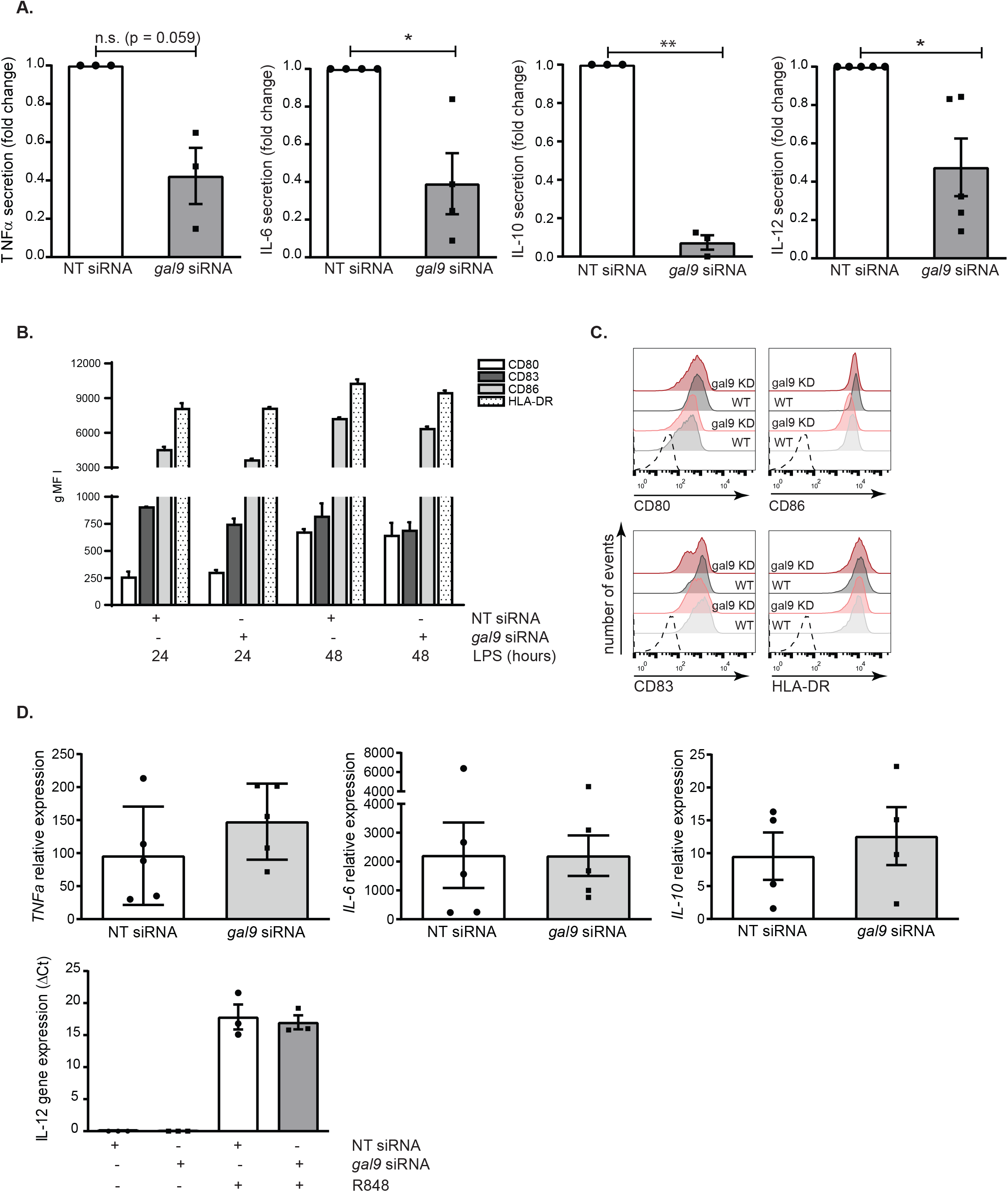
Galectin-9 is required for cytokine secretion in moDCs. **A.** Day 5 NT or *gal9* siRNA-transfected moDCs were incubated with LPS or R848 for 16 h or with zymosan for 24 h and the secretion of TNFα, IL-6, IL-12 and IL-10 determined by ELISA. Values were normalised to that of NT siRNA control cells for each individual experiment and results are expressed as fold change. All graphs represent average cytokine levels ± SEM from three to five independent donors. One column t-test was performed. **B.** NT or *gal9* siRNA-transfected moDCs were incubated with LPS for the indicated time points after which the membrane expression of CD80, CD83, CD86 and HLA-DR was determined by flow cytometry. Graph represents average membrane expression from two donors out of four analysed. gMFI = geometric mean fluorescence intensity. **C.** Representative flow cytometry plots from data shown in (B) from one donor. Dotted line represents isotype negative control, grey populations = NT siRNA moDCs; red populations = *gal9* siRNA moDCs. For both, light intensities show protein levels after 24 h LPS and dark intensities depict protein levels after 48 h LPS treatment. **D.** Day 5 NT and *gal9* siRNA transfected moDCs were treated with 1 μg/ml LPS for 3 h, zymosan (1:5 ratio) for 6 h or 4 μg/ml R848 for 8 h prior to being harvested and RNA extracted for reverse transcription. The resulting cDNA was then analysed by qRT-PCR using specific primers against TNFα, IL-6, IL-10 and IL-12. Actin was used as the reference gene for all panels. Results show mean ± SEM ΔCt values of three independent donors performed in duplicate. n.s p>0.05; * p<0.05; ** p<0.005; *** p< 0.001. For IL-12 analysis, no Ct values were detected for the untreated control.

We then performed quantitative reverse transcription (qPCR) for these cytokines to determine whether the deficiency in their secretion was due to an impairment in gene transcription or alternatively, responded to a defect in the intracellular cytokine trafficking. No differences in gene expression were found between gal-9 KD and WT cells for *IL-6, TNFα* and *IL-10* (Figure 1D). *IL-12A* gene expression levels were undetectable under basal conditions and thus only ΔCt could be calculated (Figure 1D). Overall, this data shows that galectin-9 is not involved in *de novo* transcription of cytokines in response to a stimulus, but rather in cytokine intracellular trafficking or in their secretion at the plasma membrane.

### Galectin-9 is involved in cytokine trafficking

Newly formed cytokines do not accumulate intracellularly but are continuously secreted by trafficking through the <u>e</u>ndoplasmic <u>r</u>eticulum (ER), the Golgi complex and the endosomal network to the plasma membrane (Revelo et al, 2019; Verboogen et al, 2018). Having ruled out that differences in gene transcription explained the cytokine secretion defect observed upon galectin-9 depletion, we hypothesised that galectin-9 could play a role in cytokine trafficking to the plasma membrane. To study this, we treated WT and gal-9 KD cells with LPS and measured intracellular cytokine accumulation by focusing on TNFα (Figure 2). Concomitant to an impaired cytokine secretion, TNFα was found to significantly accumulate intracellularly in gal-9 KD cells after 6 h of LPS treatment although this effect was gone at 16 h, most likely due to cytokine degradation (Figure 2A and 2B). As a positive control, ER-Golgi trafficking was inhibited using the Golgi inhibitors Brefeldin A and Monensin. Blocking cytokine trafficking caused TNFα to accumulate intracellularly in both WT and gal-9 KD cells as expected, although the intracellular levels were significantly higher in galectin-9 depleted cells (Figure 2A and 2B). To substantiate the flow cytometry data, WT or galectin-9 depleted cells were treated as before and stained against TNFα, galectin-9 and DAPI as nuclear marker. Confocal microscopy confirmed the intracellular cytokine accumulation in galectin-9 depleted cells upon LPS treatment and thus the involvement of galectin-9 in cytokine trafficking (Figure 2C and Supplementary Figure S3). In agreement to what was observed by flow cytometry, the percentage of TNFα positive DCs was markedly higher upon galectin-9 depletion (Figure 2D). The amount of TNFα accumulation, illustrated here by the integrated density, was also found to be significantly enhanced in gal-9 KD DCs compared to their WT counterparts (Figure 2E).

**Figure 2.**
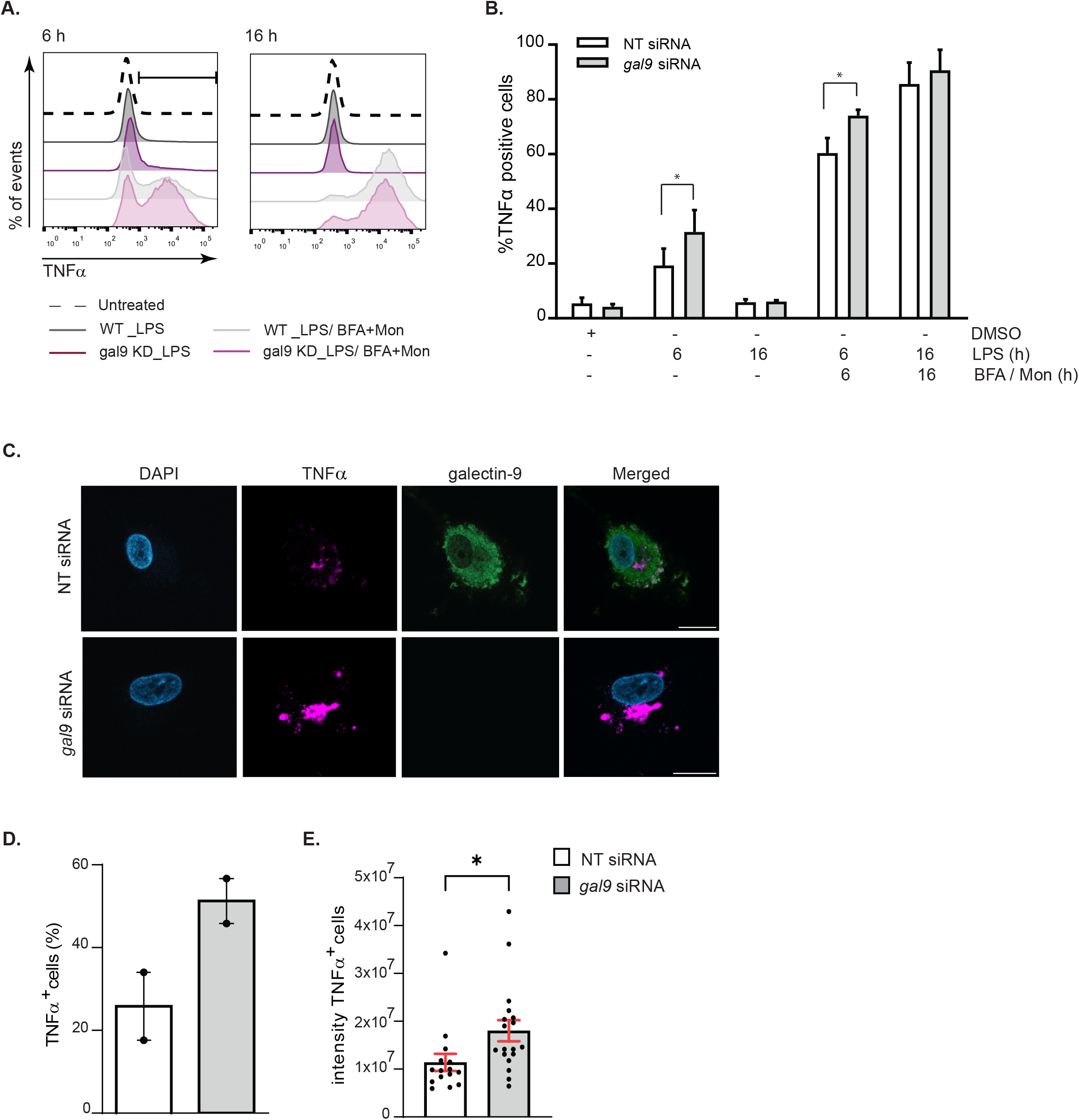
Cytokines accumulate intracellularly upon galectin-9 depletion. **A.** Day 5 NT and *gal9* siRNA transfected moDCs were treated for 6 hours with 1 μg/ml LPS alone or in combination with the Golgi inhibitors Brefeldin A (BFA, 10 μg/ml) and Monensin (Mon, 2 μM) prior to being harvested and the intracellular levels of TNFα analysed by flow cytometry. Dotted line shows TNFα levels in untreated cells. Grey depicts NT siRNA DCs and red shows *gal9* siRNA DCs (dark: cells treated with LPS (1 μg/ml) and light: cells treated with LPS + the Golgi inhibitors). **B.** Quantification and statistical analysis of experiments depicted in (A). Results show the mean value ± SEM for four independent donors. Unpaired students t-test was conducted between NT and *gal9* siRNA-transfected cells. **C.** Representative images from results shown in (A) of cells treated with LPS only and stained against TNFα, galectin-9 and DAPI for nuclear localisation. Scale bar: 10 μm. **D** and **E.** Confocal microscopy quantification of the number of TNFα positive cells (%) (D) and the integrated TNFα intensity of such cells (E) in NT and *gal9* siRNA DCs treated with LPS. Images are representative of two independent donors. Unpaired students t-test was conducted between NT and *gal9* siRNA-transfected cells. * p < 0.05.

Cytokines that fail to be secreted to the extracellular matrix need to be degraded to avoid cellular toxicity (Revelo et al, 2019). Given that the intracellular cytokine accumulation observed in gal-9 KD DCs upon LPS treatment was not sustained over time (Figure 2B), we hypothesised that galectin-9 depleted cells degraded the cytokine pool that could not be secreted. To test this, we treated WT and gal-9 KD cells with bafilomycin to block cytokine degradation *via* autophagy and measured TNFα intracellular levels using flow cytometry. As expected, inhibition of autophagy enhanced cytokine accumulation within the cytosol of DCs. This phenotype though, was markedly increased in cells depleted for galectin-9 compared to their WT counterparts as evidenced by quantifying the number of cells that showed cytokine accumulation (Figure 3A) as well as at the amount of cytokine that accumulated intracellularly (Figure 3B). Immunofluorescence performed on WT and gal-9 KD DCs treated with LPS and Bafilomycin confirmed these results (Figure 3C and supplementary Figure 4) and quantification of TNFα signal also showed the cytokine to significantly accumulate in gal-9 KD cells (Figure 3D).

**Figure 3.**
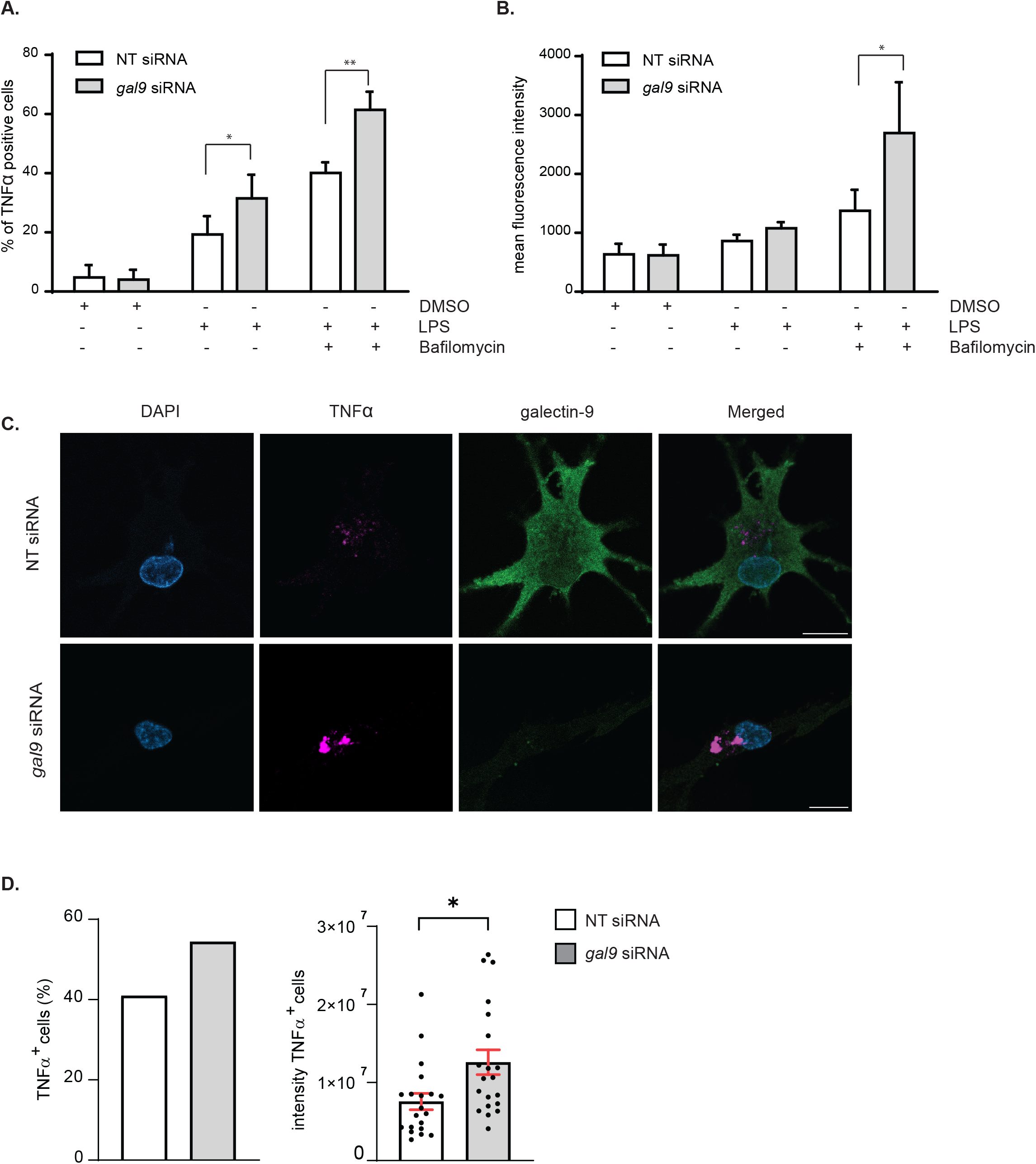
Galectin-9 depleted moDCs eliminate intracellular cytokines via lysosomes. **A** and **B.** Day 6 NT and *gal9* siRNA-transfected moDCs were treated with LPS (1 μg/ml) alone or with Bafilomycin (200 μM) for 6 hours after which cells were harvested and intracellular levels of TNFα determined by flow cytometry. Graph represents the % of TNFα positive cells (A) or the amount of TNFα present intracellularly represented by the mean fluorescence TNFα intensity (B). Graphs show mean value ± SEM of three independent donors. Paired students t-test was conducted between NT and *gal9* siRNA-transfected cells. **C.** Representative images from NT and *gal9* siRNA moDCs treated as per (A). Scale bar: 10 μm. **D.** Confocal microscopy quantification of the number of TNFα-positive cells (%) (left) and the integrated TNFα intensity of such cells (right) in NT and *gal9* siRNA DCs treated with LPS and Bafilomycin. Data represents mean ± SEM of two independent donors. Unpaired students t-test was conducted between NT and gal9 siRNA-transfected cells. * p< 0.05; ** p < 0.005.

Next, we examined the intracellular cytokine trafficking pathway in more detail to identify the organelle(s) in which cytokines accumulated *en route* to the plasma membrane for secretion in galectin-9 depleted DCs. To meet this end, fluorescence microscopy experiments were performed on WT and gal-9 KD cells treated with LPS. Upon galectin-9 depletion we observed co-localisation of TNFα at an intracellular pool that overlapped with compartments positive for the Golgi marker GM130 (Figure 4A), which was also confirmed by quantifying Manders correlation coefficient (Figure 4B). We did not detect cytokine co-localisation for those compartments positive for the early endosome marker (<u>e</u>arly <u>e</u>ndosomal <u>a</u>utoantigen 1 (EEA1)), (Figure 4C and 4D and Supplementary Figure 5A). Concomitant with flow cytometry data presented before, WT cells did not show intracellular TNFα accumulation upon LPS treatment alone at any of the cellular compartments analysed and TNFα only accumulated in the Golgi upon its blockade with Brefeldin A (supplementary Figure 5B). Nonetheless, we observed TNFα localisation in some early endosomes positive for EEA1 in WT cells, which is indication of a normal cytokine trafficking towards the plasma membrane (Figure 4C, 4D and Supplementary Figure 5A). Next, we investigated cytokine ER-Golgi trafficking in live cells upon galectin-9 depletion using the retention using selective hooks (RUSH) system, which allows for the synchronised release of TNFα-GFP protein from the ER using biotin (Boncompain et al, 2012) (Figure 5A). For this, a GFP-tagged TNFα construct was fused to an ER retention protein coupled to the streptavidin binding protein. WT and gal-9 KD DCs were transfected with the RUSH construct and live imaging was performed to visualise TNFα trafficking (Figure 5B and Supplementary movies 1 and 2). Both WT and gal-9 KD expressed the construct to similar levels and TNFα achieved maximal Golgi localisation approximately 20 min after biotin was added (Figure 5B). In agreement with our previous findings, TNFα exited the Golgi complex in WT DCs after this time but its traffic was halted in gal-9 KD cells (Figure 5B and 5C). Confocal imaging of WT and gal-9 KD moDCs transfected with the RUSH construct confirmed the accumulation of TNFα at the Golgi apparatus in galectin-9 depleted cells (Figure 5D and E), which was also quantified using the Manders correlation coefficient (Figure 5F).

**Figure 4.**
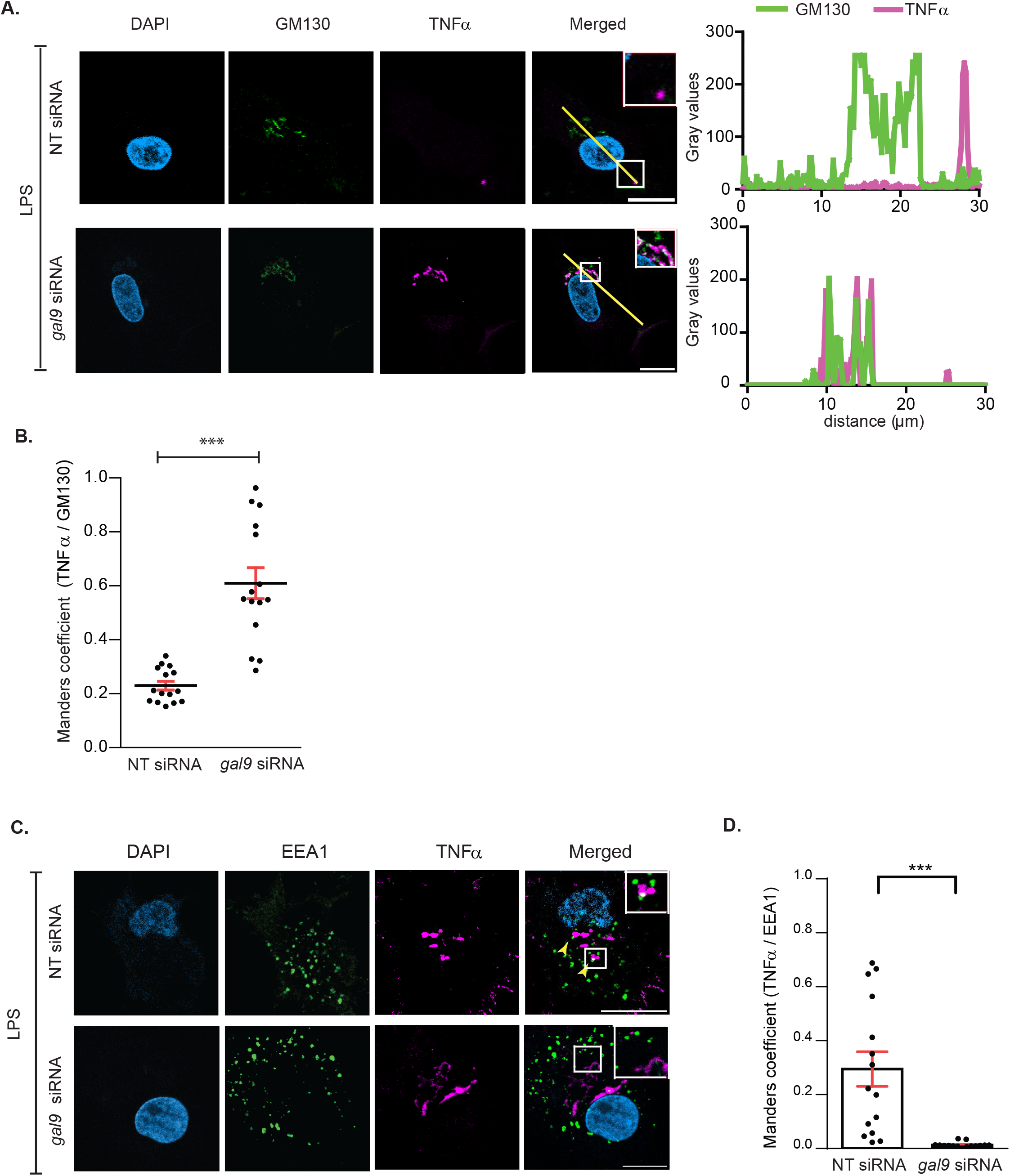
Cytokines accumulate at the Golgi apparatus in galectin-9 deficient DCs. **A.** NT and *gal9* siRNA-transfected moDCs were treated with 1 μg/ml LPS for 2 hours after which cells were fixed and stained with specific antibodies against TNFα, the Golgi marker GM130 and DAPI for nuclear staining. Representative airyscan confocal images of three independent experiments are shown. Graphs: fluorescent cross-sections as indicated. **B.** Co-localisation of TNFα and GM130 quantified by Manders correlation coefficient in moDCs treated as per (A). Fifteen cells from two representative donors are quantified. **C.** NT and *gal9* siRNA-transfected moDCs were treated with 1 μg/ml LPS for 6 hours after which cells were fixed and stained with specific antibodies against TNFα, EEA1 and DAPI for nuclear localisation. Representative airyscan confocal images from two donors are shown. Arrows indicate sites of TNFα and EEA1 colocalisation. **D.** Manders correlation coefficient was calculated for images shown in (C). Paired student t-test was conducted between NT and *gal9* siRNA transfected moDCs. *** p< 0.001.

**Figure 5.**
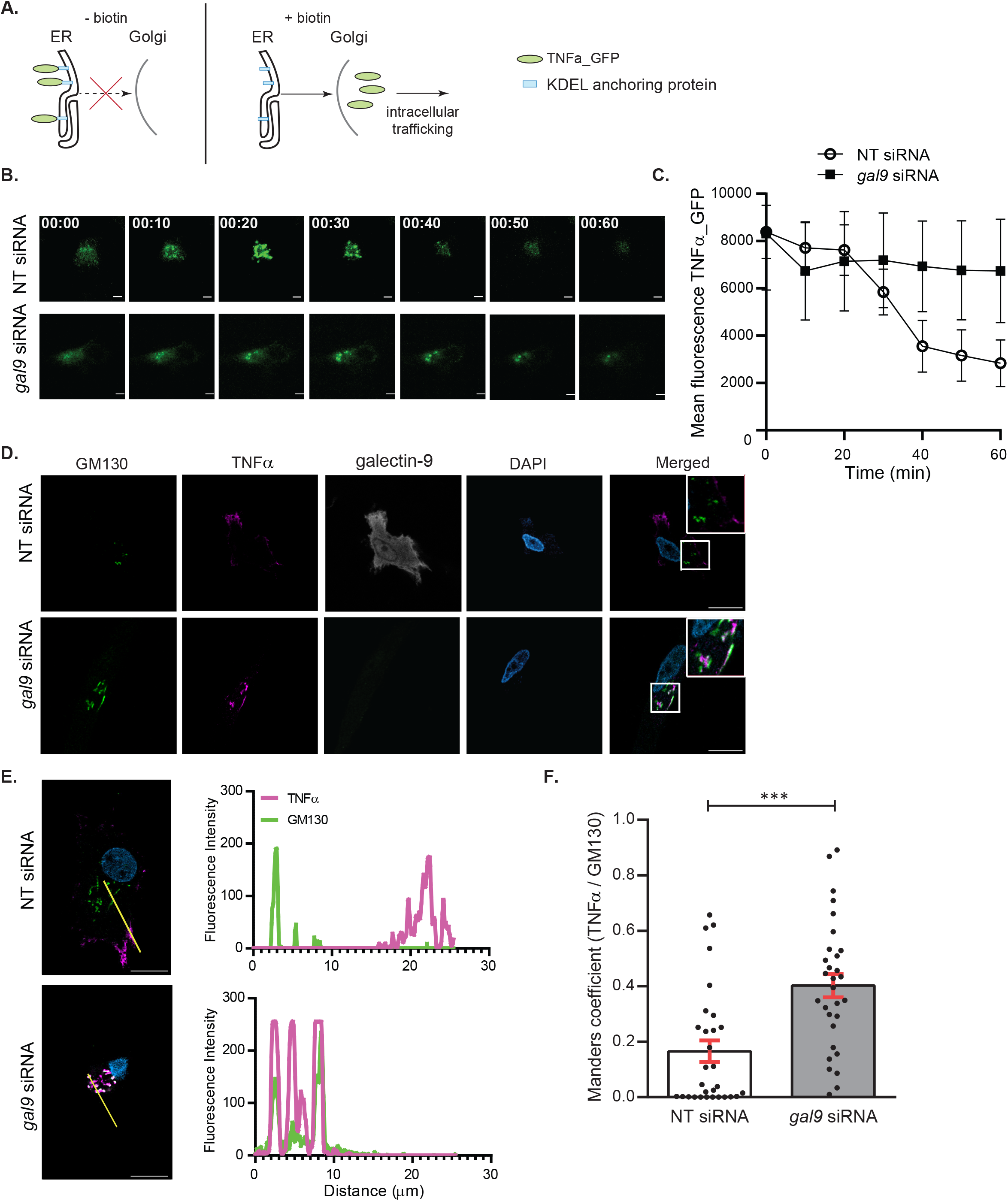
Post-Golgi cytokine trafficking is impaired upon galectin-9 depletion. **A.** Scheme depicting the experimental design based on the RUSH system. In the absence of biotin (left panel), the TNFα_GFP is trapped at the ER by the lumenal Str-KDEL hook and no cytokine trafficking post-ER can occur. When biotin is added (right panel), biotin outcompetes the interaction with streptavidin, allowing TNFα_GFP to freely traffic to the Golgi and the endosomal compartments. **B.** NT and *gal9* siRNA transfected moDCs were transfected with a GFP-labelled TNFα RUSH construct and treated with biotin to induce GFP-TNFα trafficking. Images represent snapshots of livecell imaging GFP-TNFα at the indicated time points. Scale bar: 10 μm **C.** Quantification of images shown in (B). Data represents mean intensity ± SEM from two independent donors. **D.** moDCs treated as per (B) were incubated with biotin for 30 min prior to being fixed and stained for GM130 as Golgi marker, galectin-9 and DAPI for nuclear localisation. Representative images are shown. Scale bar: 10 μm. **E.** Representative line scans of cells shown in (D) depicting GM130 (yellow) and TNFα (in red) intensity across the cell. **F.** Manders correlation coefficient was calculated for images shown in (D). Paired student’s t-test was conducted between NT and *gal9* siRNA transfected moDCs. Data represents mean correlation coefficient ± SEM of two independent donors. Fifteen cells were analysed for each condition per donor. *** p < 0.001.

Overall, these results demonstrate that galectin-9 is essential for cytokine trafficking to the plasma membrane via the endosomal machinery.

### Galectin-9 interacts with VAMP-3 to regulate cytokine trafficking

Newly synthesised cytokines are transported from the ER-Golgi complex to the plasma membrane by vesicular trafficking, a highly regulated process that involves, amongst others, proteins from the SNARE family of membrane fusion proteins (Dingjan et al, 2018). To unravel the mechanism underlying galectin-9 function in cytokine trafficking, we used an unbiased immunoprecipitation (IP)-mass spectrometry approach to identify new galectin-9 binding partners in DCs. Gal-9 was found to be significantly enriched in the Gal-9 IP compared to isotype control, confirming the quality of the pulldown experiment. Analysis of all putative galectin-9 binding partners focused on components of the vesicular trafficking identified <u>v</u>esicle-<u>a</u>ssociated <u>m</u>embrane <u>p</u>rotein 3 (Vamp-3) to interact specifically with galectin-9 (Figure 6A). Vamp-3 is a R-SNARE family member located in the early and recycling endosomal compartments (Bajno et al, 2000; Manderson et al, 2007). In exocytosis processes, it accumulates in nascent endosomes outside of the Golgi complex and it is crucial in controlling their docking and fusion to the plasma membrane for extracellular release (Barysch et al, 2009; Mallard et al, 2002). Vamp-3 is crucial for cytokine trafficking to the plasma membrane prior to their release and thus we examined if galectin-9 interaction with Vamp-3 explained the impairment in cytokine secretion in gal-9 KD cells. To confirm the molecular interaction between both proteins, co-immunoprecipitation experiments were performed in resting and in activated WT DCs. As shown, galectin-9 was found to associate with Vamp-3 both under basal conditions and upon DC activation after LPS treatment, demonstrating their molecular interaction and highlighting a novel binding partner for galectin-9 (Figure 6B). Galectin-9 interaction with Vamp-3 was also confirmed in THP-1 cells (a monocytic cell model widely used to mimic monocytes/macrophage function), suggesting their association is conserved across (immune) cells and further confirming a role for galectin-9 in intracellular trafficking in mammalian cells (Figure 6C). Further validating their relation, analysis of Vamp-3 protein levels revealed galectin-9 to specifically regulate Vamp-3 expression as Vamp-3 protein levels were diminished in gal-9 KD DCs (Figure 6D and 6E). Interestingly, depletion of galectin-9 did not alter the protein levels of other trafficking proteins (SNAP23 or Rab11a) known to be involved in cytokine trafficking (Dingjan et al, 2018). This confirms a specific functional interaction between galectin-9 and Vamp-3.

**Figure 6.**
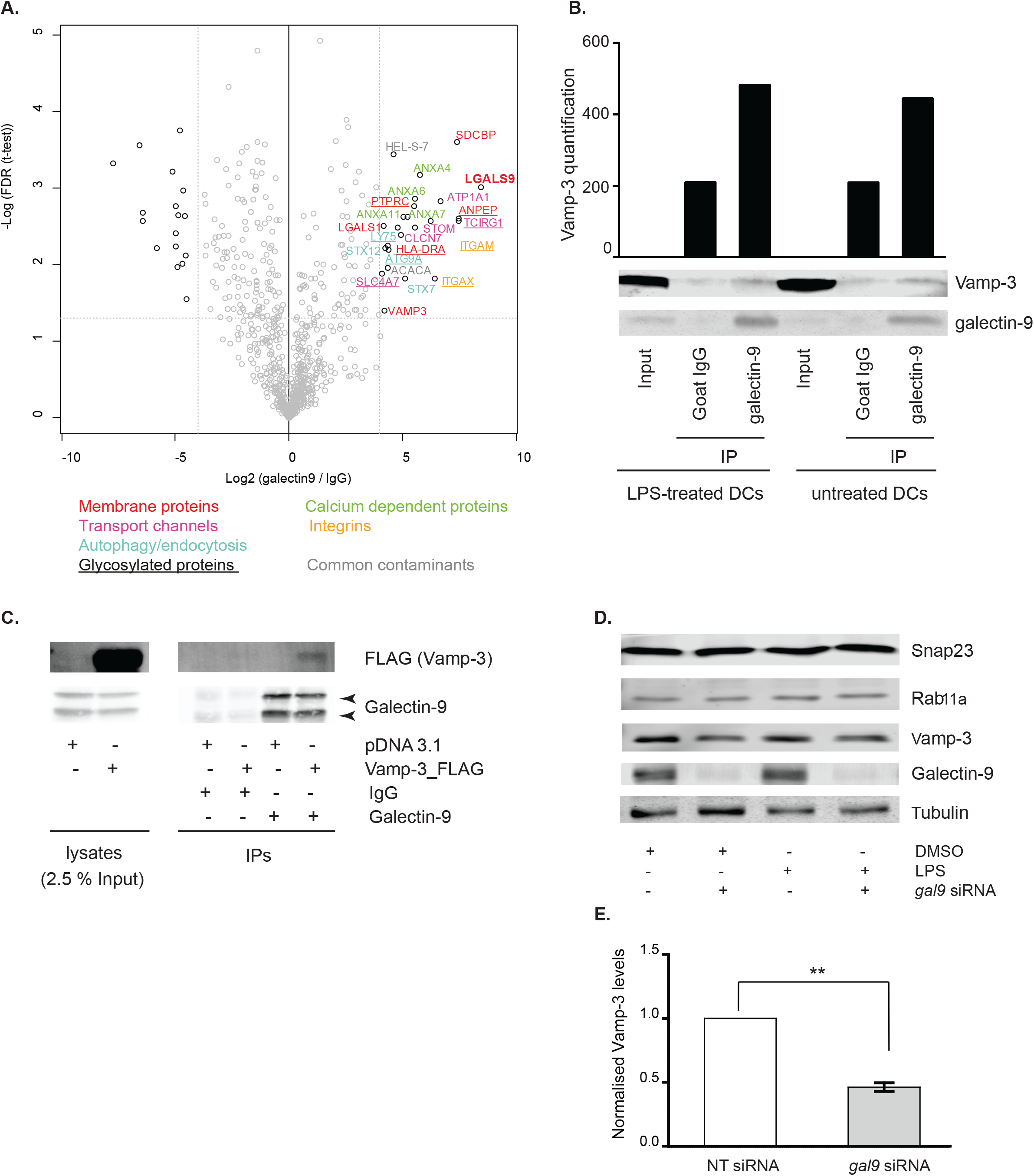
Galectin-9 interacts with Vamp-3 in DCs. **A.** moDCs were lysed and whole-cell extracts incubated with either anti-galectin-9 specific antibody or an isotype control (goat IgG). All interacting proteins were isolated and identified by mass-spectrometry. Volcano plot depicts differentially enriched galectin-9 (right) and isotype (left) interacting proteins. **B.** moDCs were prepped as per (A) but immunoprecipitated (IP) complexes were resolved and probed with galectin-9 and Vamp-3 specific antibodies. Graph shows quantification of Vamp-3 content in each sample using ImageJ. **C.** THP-1 cells were transfected with empty vector or plasmid containing 3x FLAG_Vamp-3, lysed and whole-cell extracts incubated with either anti-galectin-9 specific antibody or an isotype control (goat IgG). IP complexes were resolved and probed with galectin-9 and Vamp-3 specific antibodies. Immunoblot shown is representative of four independent experiments. **D**. NT or *gal9* siRNA transfected moDCs were treated with LPS or DMSO as negative control for 6 h prior to being lysed and whole lysates probed against the specified proteins. Tubulin was used as loading control. Immunoblot shown is representative of three independent experiments. **E.** Graph depicts Vamp-3 level in galectin-9 depleted moDCs normalised to tubulin and made relative to the corresponding NT siRNA sample. Graph shows mean ± SD of three independent donors. ** p < 0.005.

Under basal conditions, Vamp-3 is predominantly present in a cellular area juxtaposed to the nucleus (Verboogen et al, 2017). Activation of WT DCs in response to LPS treatment and consequent cytokine trafficking caused Vamp-3 to redistribute throughout the cell and towards the plasma membrane (Figure 7A). Interestingly, Vamp-3 failed to relocate upon activation in galectin-9 depleted DCs and stayed close to the nucleus (Figure 7A). Quantification of multiple cells revealed Vamp-3-containing endosomes to mostly locate within 10 μm from the nuclei under basal conditions in both WT and gal-9 KD DCs (Figure 7B). Nonetheless, already in the absence of any stimuli Vamp-3 was found to be significantly closer to the nucleus (0 - 5 μm distance) in the absence of galectin-9, which suggests galectin-9 is important for Vamp-3 localisation also under steady state conditions. Activation of the trafficking machinery resulted in Vamp-3 signal to evenly distribute across WT DCs and towards the plasma membrane for cargo release (distances of 0 - 20 μm from the nuclei, Figure 7B). Relocation of Vamp-3 was not apparent in galectin-9 depleted DCs and the majority of Vamp-3 endosomes stayed close to the nucleus (0 - 10 μm distance) upon DC activation with LPS treatment (Figure 7B). Immunofluorescence experiments confirmed Vamp-3 to be retained within the Golgi complex upon LPS treatment in Gal-9 KD DCs as Vamp-3 signal accumulated within areas positive for the Golgi marker GM130 (Figure 7C). Vamp-3 did not co-localise with the ER marker PDI in either WT or gal-9 KD cells suggesting galectin-9 is not involved in the ER-Golgi trafficking but rather controls post-Golgi transport into the endosomal compartment (Figure 7C). Relocation of Vamp-3 after LPS activation was apparent in WT cells as Vamp-3 co-localised with compartments positive for EEA1 (Figure 7C).

**Figure 7.**
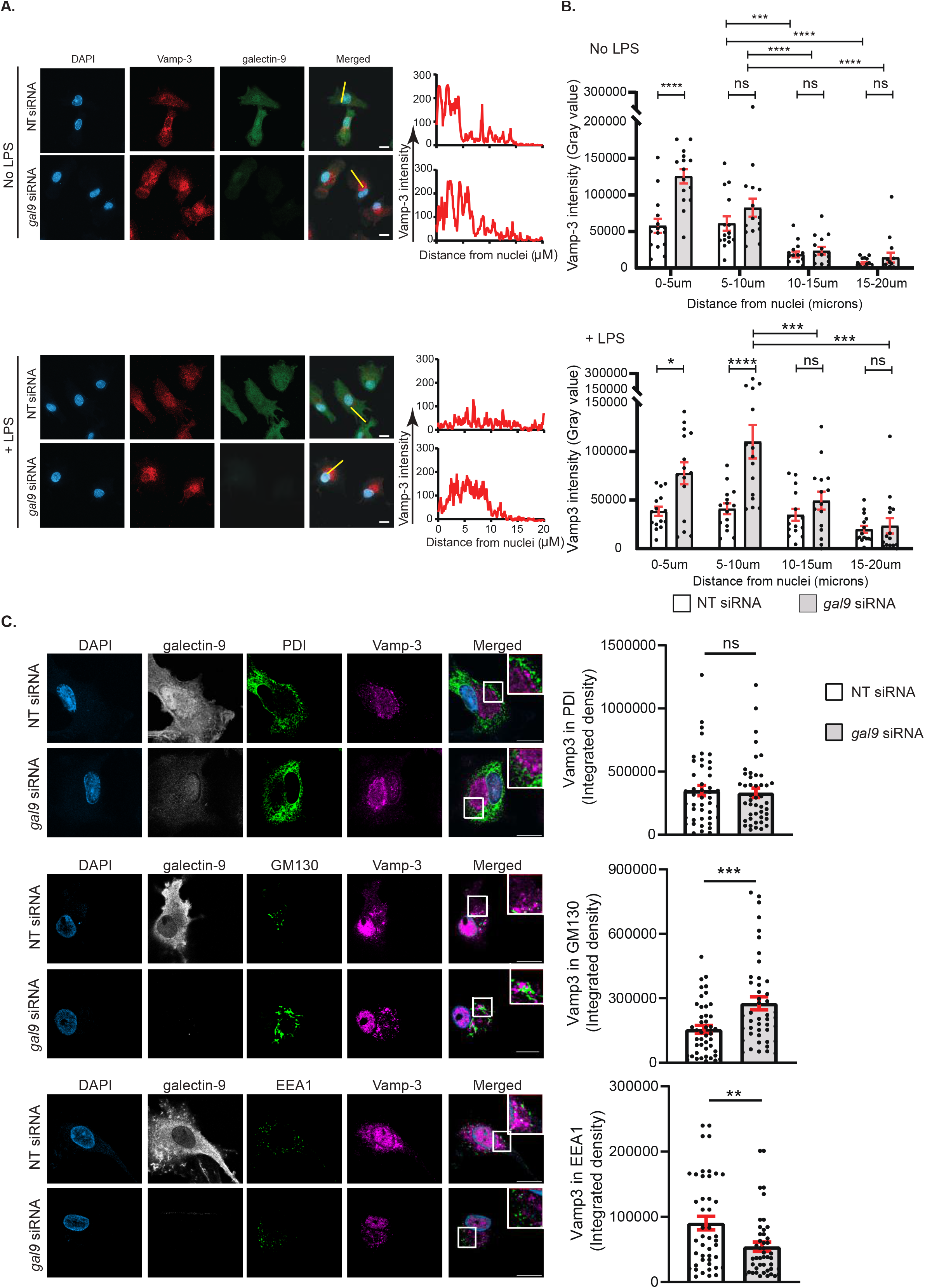
Galectin-9 regulates Vamp-3 intracellular localisation. **A.** NT and *gal9* siRNA transfected moDCs were incubated with LPS for 4 h prior to being fixed in methanol and stained for Vamp-3, galectin-9 and DAPI for nuclear localisation. Representative images for each treatment are shown. Graphs: fluorescent cross-sections as indicated depicting Vamp-3 intensity across the cell. Scale bar: 10 μm. **B.** Line scan quantification of cells shown in (A). Data represents mean Vamp-3 intensity for the specified distance from the nuclei ± SEM of one representative donor out of three. Fifteen cells were analysed per condition. Two-way ANOVA was performed for statistical analysis. **C.** Representative images of DCs treated as in (A), fixed in 4 % PFA and stained against galectin-9, Vamp-3, PDI, GM130, EEA1 and DAPI for nuclear localisation. Graph depicts Vamp-3 integrated density values for three independent donors ± SEM. Fifteen cells were analysed per donor. Unpaired t-test was conducted between NT and *gal9* siRNA samples. Please note that PFA-fixed cells also show Vamp-3 nuclear staining as a result of the fixation process. n.s. p > 0.05; ** p < 0.01; *** p<0.001; **** p< 0.0001.

Taken together, our data demonstrates that intracellular galectin-9 controls cytokine trafficking *via* its functional interaction with Vamp-3, which underlies Galectin-9 requirement to direct Vamp-3 containing endosomes to the plasma membrane for cytokine secretion in DCs.

## Discussion

Vesicle trafficking is pivotal to maintain cellular homeostasis and coordinate exo- and endocytosis. Intracellular protein transport spans over different compartments (ER, Golgi complex and endosomes) that form a dynamic network in which vesicles deliver cargo from one to the other. This process is fundamental to ensure the timely delivery of specific components to the plasma membrane, for instance during (immune) cell activation. Illustrating this, DC stimulation markedly enhances the intracellular trafficking machinery required to transport specific surface receptors to the plasma membrane as well as the secretion of (immune)modulatory cytokines. In this study, we identify galectin-9, a tandem repeat member of the galectin family, as a key mediator of intracellular cytokine trafficking and report a novel functional interaction with the SNARE protein Vamp-3. Vamp-3 locates in early/recycling endosomes and is crucial for cytokine trafficking to the plasma membrane prior to their release (Manderson et al, 2007; Murray et al, 2005). Depletion of galectin-9 results in lower Vamp-3 protein levels as well as in its aberrant subcellular localisation, which prevents cytokines from entering the endosomal transport machinery and causes their accumulation at the Golgi. Thus, this study shows that galectin-9 interacts with Vamp-3 and governs its redistribution throughout the cytosol upon DC activation, in turn allowing for Vamp-3 mediated trafficking to occur (Figure 8).

**Figure 8.**
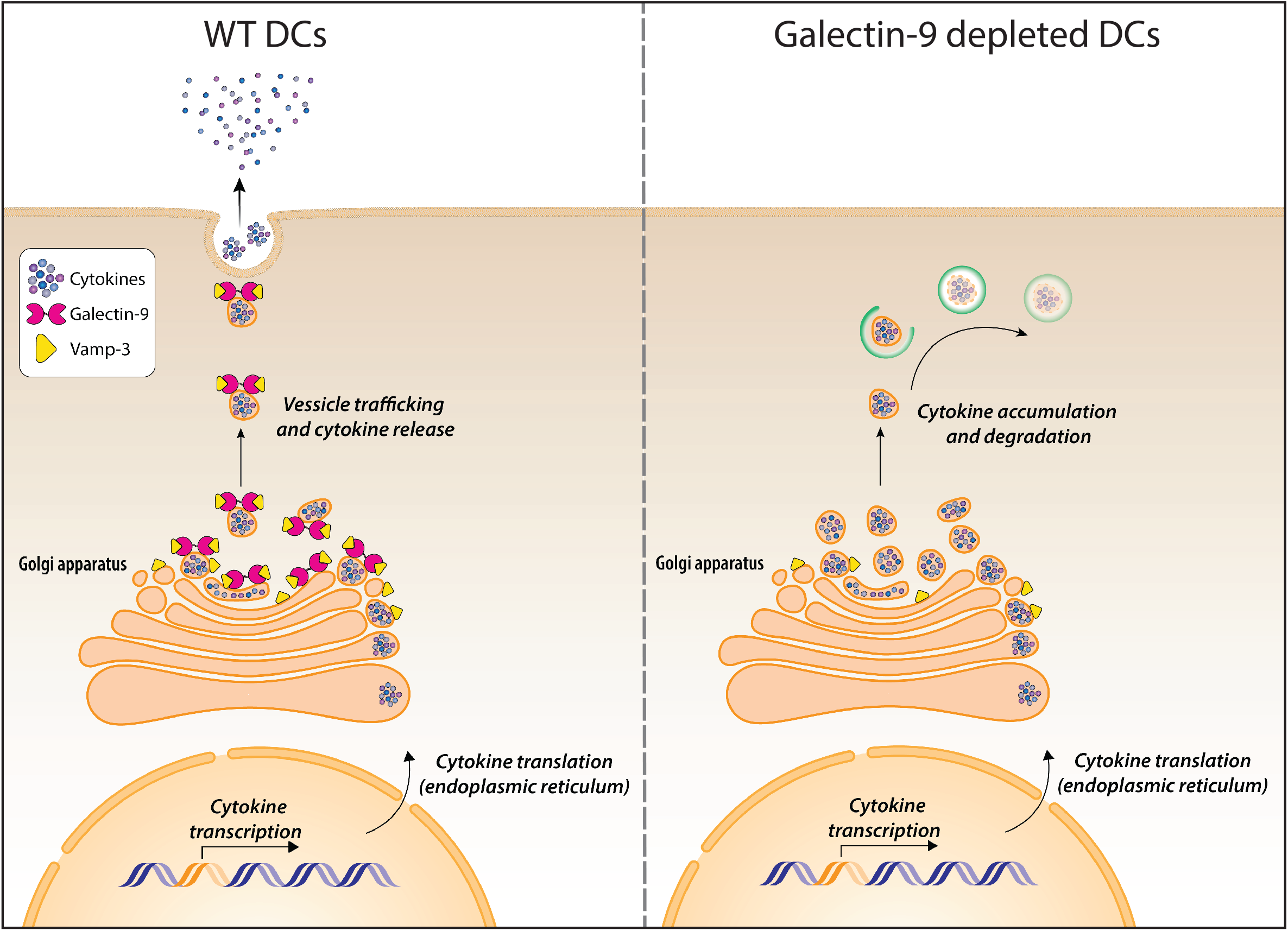
Graphical abstract. Galectin-9 interacts with Vamp-3 to regulate cytokine secretion in DCs. Upon stimuli, DCs activate and synthesise *de novo* cytokines that need to be transported *via* the ER/Golgi complex to the endosomal compartment, from where they will traffic to the plasma membrane and be released to the extracellular matrix. Vamp-3 is a SNARE protein crucial in facilitating cytokine secretion of the Golgi complex to the plasma membrane in DCs. Galectin-9 interacts with Vamp-3 and facilitates Vamp-3 redistribution from the edge of the Golgi complex to the rest of the cytoplasm upon DC activation. In the absence of Galectin-9, Vamp-3 is sequestered at the Golgi compartment, which results in the accumulation of cytokine-containing vesicles in the Golgi complex that eventually undergo lysosomal degradation.

Galectins are a family of β-galactoside binding proteins which have been identified as novel membrane organisers (Elola et al, 2015; Lajoie et al, 2009; Nabi et al, 2015). They are also abundantly expressed intracellularly and we and others have reported several members of the galectin family to modulate (immune) cell function *via* carbohydrate-independent interactions. In particular, galectin-9 is known to influence various processes such as cell adhesion, aggregation or apoptosis and we have previously identified a role for galectin-9 in phagocytosis by DCs (John & Mishra, 2016; Querol Cano et al, 2019). In contrast, very little is known regarding the involvement of any member of the galectin family in cytokine intracellular trafficking or secretion. The currently available data on this topic mainly refers to either descriptive observations or to the correlation of specific cytokines and galectin protein levels, while the regulatory mechanisms are largely lacking (Gordon-Alonso et al, 2018). We now demonstrate a role for endogenous galectin-9 in mediating cytokine secretion in DCs via its function in regulating vesicle trafficking between the Golgi network and the plasma membrane. Our work is in line with studies reporting that extracellular recombinant galectin-9 enhances IL-6 and IL-8 secretion in human mast cells (Kojima et al, 2014) and IL-12 secretion in moDCs (Dai et al, 2005). However, underlying molecular mechanisms have not been resolved and the reported effects could be carbohydrate dependent (mast cells) or carbohydrate-independent (DCs), suggesting that galectin-9-mediated pathways differ in distinct cell types. These different results could also be explained by the use in these studies of excess recombinant galectin-9, which will bind to unknown carbohydrate-containing membrane proteins at the cell surface and not necessarily alter the intracellular cytokine secretion machinery. To decipher how endogenous galectin-9 protein functions, we exploited knockdown technology in primary human cells. Our data specifically demonstrates the intracellular interaction between endogenous Vamp-3 and galectin-9 and that galectin-9 depletion results in defective TNFα trafficking from the ER-Golgi to the cell surface. Still, it cannot be excluded that membrane bound galectin-9 may (indirectly) also participate in controlling cytokine secretion *via* other unknown mechanisms. Furthermore, and highlighting the translational relevance of our findings, galectin-9 plasma levels have been shown to correlate with elevated cytokines in COVID-19 patients and addition of recombinant galectin-9 induced IL-6 and TNFα secretion in those patients (Bozorgmehr et al, 2021).

Our ELISA data revealed a significant decrease in cytokine release upon galectin-9 depletion, whilst maintaining cytokine *de novo* synthesis. Concomitantly, intracellular TNFα staining confirmed our hypothesis that if cytokine production was preserved between WT and gal-9 KD cells but secretion was impaired in the latter, cytokines should accumulate in the cytosol of galectin-9 depleted cells. Furthermore, we have demonstrated that galectin-9 deficient DCs eliminate the excess of cytosolic cytokines that cannot be released onto the extracellular matrix *via* the lysosomal machinery. This is in agreement with published literature that reported the importance of autophagy as a degradation mechanism in cytokine regulation (Jiang et al, 2019; Wu et al, 2016).

Galectin-9 did not alter the trafficking of other proteins to the plasma membrane upon DC activation, suggesting this lectin is not involved in global intracellular trafficking processes. In agreement with our data, galectin-9 has not been reported to participate in protein trafficking although other members of the galectin family have been described to aid in the cytosolic transport of various cell surface components in epithelial cells and erythrocytes (Barres et al, 2010; Delacour et al, 2006; Stechly et al, 2009; Straube et al, 2013). The apparent dichotomy we here report between the normal trafficking of membrane components and the defect in cytokine secretion in gal-9 KD DCs can be explained by the different routes by which these proteins arrive to the plasma membrane. For instance, and although activation-induced cell surface display of several maturation markers depends on *de novo* synthesis, it has also been described that CD83 and MHC-II reside in distinct pools of recycling endosomes that fuse to the plasma membrane upon DC activation (Klein et al, 2005). Similarly, ubiquitination serves as an important mechanism by which DCs regulate CD86 expression upon activation (Baravalle et al, 2011).

Cytokine (constitutive) release in DCs is primarily regulated at the transcriptional/translational level after a stimulus triggers their biosynthesis (Chiaruttini et al, 2016; Tanaka et al, 2016). This is in agreement with our own data, which demonstrates a rapid upregulation in cytokine gene transcription upon DC activation. Cytokine transcription was not impaired in galectin-9 depleted DCs demonstrating galectin-9 is not involved in the activation of the translational machinery in response to inflammatory stimuli. Newly formed cytokines are then subsequently trafficked through the endoplasmic reticulum, the Golgi complex and the vesicle network to the plasma membrane (Stanley & Lacy, 2010; Verboogen et al, 2018). In macrophages and DCs, newly synthesised cytokines do not traffic directly from the Golgi apparatus to the plasma membrane, but are instead routed *via* recycling endosomes that partially overlap with Vamp-3 (early/recycling endosomes) and the transferrin receptor (TfR), present in recycling endosomes (Manderson et al, 2007; Verboogen et al, 2018). In agreement with galectin-9 governing Vamp-3 function in transporting cytokines away from the Golgi complex, our microscopy analysis revealed TNFα to accumulate at the Golgi after 4-6 h of LPS treatment in gal-9 KD cells. This indicates cytokine trafficking is abrogated after this point upon galectin-9 depletion and cytokines cannot be further trafficked, thus galectin-9 is required for cytokine transport from the Golgi to the plasma membrane. On the contrary, no cytokine could be detected at the Golgi complex of LPS-treated WT cells, which aligns with previous publications showing that in DCs and macrophages cytokine accumulation at the Golgi cannot be observed 2-4 h after LPS stimulation (Chiaruttini et al, 2016; Manderson et al, 2007; Shurety et al, 2000).

Mounting evidence suggests that post-Golgi trafficking is the rate-limiting step for the secretion of cytokines such as IL-6, IL-12 and TNFα as DCs need to upregulate their trafficking machinery to cope with the demand for cytokine production in response to stimuli (Revelo et al, 2019). In agreement, TNFα release at the plasma membrane depends on the protein levels of several SNAREs (i.e Vamp-3 or syntaxin-4) and knockdown of Vamp-3 has been shown to reduce IL-6 and TNFα secretion in DCs (Boddul et al, 2014; Collins et al, 2014; Mori et al, 2011). This phenotype aligns with the impairment in cytokine secretion upon galectin-9 depletion we here report and argues the functional interaction of both proteins is required for effective cytokine trafficking and release. Vamp-3 is known to interact with various syntaxin proteins at the plasma membrane, namely syntaxin-4, which allows Vamp-3 containing endosomes to fuse with the plasma membrane (Dingjan et al, 2018; Verboogen et al, 2017). We have no evidence that galectin-9 is involved in the anchoring of endosomes to the plasma membrane prior to cargo release. In line with this, our Vamp-3 microscopy data unequivocally identified an earlier impairment in the trafficking machinery of galectin-9-depleted cells, namely the inability of Vamp-3 to relocate from the Golgi complex to the cell surface. Endosomes do not randomly diffuse throughout the cytosol but are rather captured by the cytoskeleton, which provides a functional structure for vesicle transport with motor proteins that transports them along the actin and microtubule fibres (Granger et al, 2014; Klann et al, 2012). We have previously found galectin-9 to locate within the cortical actin and to actively participate in actin polymerisation during phagosome formation and thus it is possible that galectin-9 impairs Vamp-3 intracellular movement due to its interactions with the cortical actin cytoskeleton (Querol Cano et al, 2019).

In summary, our work demonstrates that galectin-9 controls cytokine intracellular trafficking and secretion through its functional interaction with the SNARE protein Vamp-3, which allows galectin-9 to govern Vamp-3 cytosolic redistribution upon cell activation, in turn allowing for Vamp-3 mediated trafficking to occur. Within the subcellular vesicle transport route, galectin-9 is required for vesicle transport from the Golgi to the plasma membrane. To the best of our knowledge, no intracellular binding partners of galectin-9 involved in the trafficking machinery have been previously reported. Given the myriad of cellular functions that depend on intracellular trafficking processes in all cells of our body, this novel mechanism reverberates in several fields within the cell biology research area. Furthermore, our findings increase our understanding of the multiple roles galectins play in governing (immune) cell function, which may also be conserved in evolution.

## Materials and Methods

### Generation of monocyte-derived dendritic cells

Dendritic cells were derived from peripheral blood monocytes isolated from a buffy coat (Sanquin, Nijmegen, The Netherlands) (de Vries et al, 2002). Monocytes isolated from healthy blood donors (informed consent obtained) were cultured for up to five days in RPMI 1640 medium (Life Technologies, Bleiswijk, Netherlands) containing 10 % foetal bovine serum (FBS, Greiner Bio-one, Alphen aan den Rijn, Netherlands), 1 mM ultra-glutamine (BioWhittaker), antibiotics (100 U/ml penicillin, 100 μg/ml streptomycin and 0.25 μg/ml amphotericin B, Life Technologies), IL-4 (500 U/ml) and GM-CSF (800 U/ml) in a humidified, 5 % CO_2_. On day 3, moDCs were supplemented with new IL-4 (300 U/ml) and GM-CSF (450 U/ml).

### Small interfering RNA knockdown

On day 3 of DC differentiation, cells were harvested and subjected to electroporation. Three custom stealth small interfering RNA (siRNA) were used to silence galectin-9 (LGALS9HSS142807, LGALS9HSS142808 and LGALS9HSS142809) (Invitrogen). Equal amounts of the siRNA ON-TARGETplus non-targeting (NT) siRNA#1 (Thermo Scientific) were used as control. Cells were washed twice in PBS and once in OptiMEM without phenol red (Invitrogen). A total of 15 μg siRNA (5 μg from each siRNA) was transferred to a 4-mm cuvette (Bio-Rad) and 5-10×10^6^ DCs were added in 200 μl OptiMEM and incubated for 3 min before being pulsed with an exponential decay pulse at 300 V, 150 mF, in a Genepulser Xcell (Bio-Rad, Veenendaal, Netherlands), as previously described (Querol Cano et al, 2019). Immediately after electroporation, cells were transferred to preheated (37 °C) phenol red–free RPMI 1640 culture medium supplemented with 1 % ultraglutamine, 10 % (v/v) FCS, IL-4 (300 U/ml), and GM-CSF (450 U/ml) and seeded at a final density of 5×10^5^ cells/ml.

### Enzyme-linked immunosorbent assay (ELISA)

Day 5 moDCs were seeded in 96 well plates at a concentration of 1×10^5 cells/well in RPMI 1640 culture medium supplemented with 1 % ultraglutamine, 10 % (v/v) FCS, IL-4 (300 U/ml), and GM-CSF (450 U/ml). Cells were then treated for 6, 16 or 24 hours with LPS (Sigma-Aldrich; 1 μg/ml), zymosan (life technologies, added at a 1:5 ratio) or R848 (Sigma-Aldrich; 4 μg/ml). After this time, cells were spun down for 2 min at 1500 rpm and supernatants collected for ELISA. TNFα and IL-6 were analysed from samples stimulated with LPS, IL-10 from samples treated with zymosan and IL-12 was measured on samples treated with R848.

Secreted levels of TNFα, IL-6 and IL-10 were analysed using human ELISA Ready-SET-Go! Kits (eBioscience) and following manufacturer’s instructions. To determine IL-12 secretion, 96-well ELISA plates were coated with coating buffer (Na_2_CO_3_ 0.1 M pH 9.6) containing IL-12p70 coating antibody (Invitrogen, clone 20C2 M122 and used at 3 μg/ml) overnight and at 4 °C. Samples were afterwards incubated for 1 hour at room temperature before adding the detection antibody (Pierce Endogen, clone M121B at 250 ng/ml). After this time, streptavidin-HRP (Invitrogen, used at 250 ng/ml) was added and samples were further incubated for 30 min at room temperature. Tetrametilbenzidine (TMB) was used to start the colorimetric reaction and when appropriate, H3PO4 was used to stop the reaction. All plates were read using an iMark Microplate absorbance reader (Bio-Rad) and processed using GraphPad Prism 6 software.

### Flow cytometry

To determine depletion of galectin-9 following siRNA transfection, single cell suspensions were stained with a goat anti-galectin-9 antibody (AF2045, R&D systems) at 40 μg/ml or isotype control as negative control for 30 min at 4 °C. Before staining, moDCs cells were incubated with 2 % human serum for 10 min on ice to block non-specific interaction of the antibodies with FcRs. A donkey-anti goat secondary antibody conjugated to Alexa Fluor 488 was used (Invitrogen; 1:400 (v/v). All antibody incubations were performed in PBA containing 2 % human serum.

When necessary, day 6 moDCs were seeded in low adhesion plates and treated with Brefeldin A (BFA; 10 ug/ml) and Monensin (eBioscience; 2 μM) for 6 or 16 hours alone or in combination with LPS (1 μg/ml). For experiments with lysosomal inhibitors, day 6 moDCs were treated with 200 nM Bafilomycin A1 (Sigma-Aldrich) alone or in combination with LPS (1 μg/ml) for 6 hours prior to being collected. Cells were then stained with the Zombi violet viability dye (BioLegend, 1:2000 dilution) prior to being fixed and permeabilised using the Fixation/permeabilisation solution kit (BD Bioscience) and following manufacturer’s instructions. Cell suspensions were then stained using the directly-labelled primary antibody TNFα_PE (BioLegend; 502929) or mouse IgG1 (eBiosciences, #12-4714-42) as negative control.

Cells were analysed with a FACSVerse or a FACSLyric instrument (BD Biosciences) and results analysed using FlowJo version X software (Tree Star, Ashland, Oregon).

### Co-immunoprecipitation and mass-spectrometry

Day 6 moDCs (10×10^6^) were detached using cold PBS, collected and lysed in 1 ml lysis buffer containing 150 mM NaCl, 10 mM Tris-HCl (pH 7.5), 2 mM MgCl_2_, 1 % Brij97, 2 mM CaCl_2_, 5 mM NaF, 1 mM NaVO_4_ and 1 mM PMSF for 30 min on ice. Cell lysates were then incubated with 2 μg of anti-galectin-9 (AF2545, R&D systems) or isotype control under rotation. After incubating for 1 h at 4 °C, agarose beads were added and samples were further incubated for 3 h. Afterwards, beads were washed five times in washing buffer (150 mM NaCl, 10 mM Tris-HCl (pH 7.5), 2 mM MgCl_2_, 0.1 % Brij97, 2 mM CaCl_2_, 5 mM NaF, 1 mM Na_3_VO_4_ and 1 mM PMSF) followed by three washes with PBS and bound proteins eluted in SDS sample buffer (62.5 mM Tris pH 6.8, 2 % SDS, 10 % glycerol). Proteins were separated by PAGE and blotted onto PVDF membranes. Membranes were blocked in TBS containing 3 % BSA and 1 % skim milk powder at room temperature for 1 h prior to be stained with specific antibodies against Vamp-3 and Galectin-9. Antibody signals were detected with HRP coupled secondary antibodies and developed using Odyssey CLx (Li-Cor) following manufacturer’s instructions. Images were retrieved using the Image Studio Lite 5.0 software.

For mass spectrometry analysis, immunoprecipitation on was performed as above. After removal of all supernatant, on-bead digestion was performed as described (Baymaz et al, 2014). In short, proteins were denatured and reduced in 50 μl of 2 M urea, 100 mM Tris-HCl pH 8.0 and 10 mM DTT. Cysteines were alkylated using 50 mM Iodoacetamide for 15 min at room temperature and proteins digested using 0.25 μg MS-grade trypsin for 2 hours at room temperature in a thermoshaker. Peptide-containing supernatants were collected and incubated with 0.1 μg fresh trypsin overnight. The next day, peptides were desalted using STAGE-tips. Samples were eluted from the stagetips using buffer B (80 % acetonitrile, 0.1 % formic acid), after which acetonitrile was evaporated using a speedvac. Samples were diluted in buffer A (0.1 % formic acid) to 10 μl, of which half was injected. Peptides were separated on a reverse phase column connected to an Easy-nLC1000 (ThermoFisher) coupled on-line to a Thermo Orbitrap Exploris 480. A buffer B gradient (30 to 95 %) was run over a period of 60 minutes, during which a dynamic exclusion list of 30 seconds was applied. RAW data was analysed using Maxquant version 1.6.0.1 (Cox & Mann, 2008), with a Uniprot database of the human proteome downloaded in June 2017. Options for match-between runs and LFQ (label free quantification) were enabled. Data was further analysed in Perseus (Tyanova et al, 2016) to filter out contaminants and reverse hits. Proteins with less than 2 peptides or less than 6 valid values were also removed. A two samples test was performed and a volcano plot was generated using R.

For immunoprecipitations performed on THP-1 cells were first transfected with 2 μg of empty vector pCDNA 3.1 or plasmid containing 3x FLAG_Vamp-3 using the Amaxa 4D nucleoporator system (Lonza) with the Amaxa SG Cell line kit and program FF-100 and following manufacturer’s instructions. The human Vamp-3 was obtained via RT-PCR from human cDNA derived from monocyte derived dendritic cells. This hVAMP3 was cloned into pCDNA3.1(+) vector containing a 3x FLAG-tag via standard molecular techniques using Hind III/ BamH I sites. The construct sequence was verified by the Radboudumc sequencing facility.

Forty-eight hours after transfection, cells were collected and lysed in 1 ml lysis buffer containing 150 mM NaCl, 25 mM Tris-HCl (pH 7.5), 2 mM EDTA, 1 % Brij97 (v/v) and phosphatase and proteins inhibitors (Roche, # 4906837001 and # 4693132001) for 30 min under end over end rotation at 4 °C. Cell lysates were then spun for 10 min at 6000 rpm at 4 °C and pre-cleared by incubating them under rotation with 25 μl of Sepharose CL-4B beads (G-Biosciences, #786-1560) for further 30 min at 4 °C. Lysates were spun as before and the supernatant transferred to a new tube from which an aliquot was taken (as input control). The remaining volume was divided into two tubes and incubated on ice with either 1 μg of anti-galectin-9 (AF2545, R&D systems) or goat isotype control (Jackson Immunoresearch, #005-000-003). After incubating for 30 at 4 °C, 25 μl of protein G sepharose beads (Cytiva, #17061801) were added and samples were further incubated for 3 h under rotation at 4 °C. Afterwards, beads were washed five times in washing buffer (150 mM NaCl, 25 mM Tris-HCl (pH 7.5), 2 mM EDTA, 1 % Brij97) and bound proteins eluted in SDS sample buffer (62.5 mM Tris pH 6.8, 2 % SDS, 10 % glycerol). Proteins were separated by SDS-PAGE and blotted onto PVDF membranes. Membranes were blocked in TBS containing 3 % BSA and 1 % skim milk powder at room temperature for 1 h prior to be stained with specific antibodies against FLAG and Galectin-9. Antibody signals were detected with HRP coupled secondary antibodies and developed using Odyssey CLx (Li-Cor) following manufacturer’s instructions. Images were retrieved using the Image Studio Lite 5.0 software.

### Western Blot

Day 6 moDCs were lysed in lysis buffer for 30 min on ice prior to being spun down at 10000 rpm for 5 min. The BCA protein assay (Pierce, ThermoFisher scientific) was conducted to determine protein concentration and following manufacturer’s instructions and for each sample, 20 μg of total protein were diluted using SDS sample buffer (62.5 mM Tris pH 6.8, 2 % SDS, 10 % glycerol).

Proteins were separated by PAGE and blotted onto PVDF membranes. Membranes were blocked in TBS containing 3 % BSA and 1 % skimmed milk powder at room temperature for 1 h prior to be stained with specific antibodies. Antibody signals were detected with HRP coupled secondary antibodies and developed using Odyssey CLx (Li-Cor) following manufacturer’s instructions. Images were retrieved using the Image Studio Lite 5.0 software. The following primary antibodies were used for Western Blotting: goat anti-Galectin-9 (AF2045, R&D systems, Minneapolis, Minnesota) at 1:1000 (v/v), anti-Vamp-3 (ab5789, Abcam) at 1:1000 (v/v), anti-SNAP23 (111203, Synaptic Systems) at 1:1000 (v/v), anti-Rab11a (sc-166523, Santa Cruz Biotechnology) at 1:1000 (v/v), mouse anti-FLAG at 1:1000 (v/v) (Sigma-Aldrich M2 clone, B3111) and rat anti-tubulin (Novus Biological, Abingdon, United Kingdom) at 1:2000 (v/v). The following secondary antibodies were used: donkey anti-goat IRDye 680 (920-32224, Li-Cor, Lincoln, Nebraska), goat anti-mouse IRDye 800 (926-32210), donkey anti-rabbit IRDye 800 (926-32213, Li-Cor), donkey anti rabbit IRDye 680 (926-68073, Li-Cor), goat anti rabbit IRDye 800 (926-32211, LiCor), goat anti-rat IRDye 680 (A21096, Invitrogen, Landsmeer, Netherlands). All secondary antibodies were used at 1:5000 (v/v).

### RNA preparation and quantitative real-time PCR

Day 6 moDCs that had been previously transfected with either NT or *gal9* siRNA were treated with either 1 μg/ml LPS for 4 h to analyse IL-6 and TNFα expression, with 1:5 ratio of Zymosan particles for 6h to study IL-10 gene expression or with 4 μg/ml R848 for 8 h to study IL-12 expression prior to being harvested, total RNA extracted using the quick-RNA miniprep kit (#R1055; Zymo research) and 1 μg reverse transcribed using the MMLV reverse transcriptase kit (#28025021; ThermoFisher scientific). *TNFα, IL-6, IL-12A* and *IL-10* gene expression was assessed by quantitative real-time PCR using FAST Reaction SYBR-Green mixture (applied Biosystems). Data was normalised to *actin* expression. Primer sequences are shown in Table 1.

**Supplementary Table 1.**
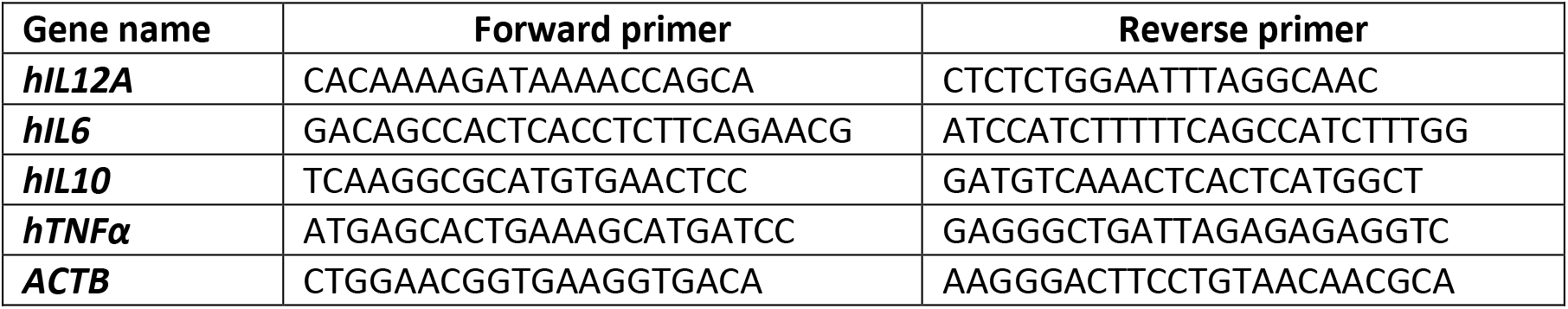
Primer sequences

### Live imaging

Day 5 moDCs previously transfected with either NT or *gal9* siRNA were transfected with 2 ug of the Str-KDEL_TNFα_SBP_EGFP plasmid (Addgene, #65278) (Boncompain et al, 2012) using the Neon transfection system (Invitrogen) and following manufacturer’s instructions. After transfection cells were seeded in DMEM (Gibco, #31966-021) on low attachment plates (Corning, #3471) and treated with 1 μg/ml LPs 3 h after transfection. Sixteen hours later, GFP positive cells were sorted out using the BD FACSMelody cell sorter (BD Biosciences) and immediately seeded onto glass bottom imaging chambers (Greiner Bio-one #627870) on DMEM to attach. Three hours later, cells were washed once in PBS and incubated in Leibovitz’s medium (Gibco; 21083-027) prior to imaging. Cells were imaged using a Leica DMI6000 epifluorescence microscope fitted with a 63x 1.4 NA oil immersion objective, a metal halide EL6000 lamp for excitation, a DFC365FX CCD camera and GFP filter sets (all from Leica, Wetzlar, Germany). Focus was kept stable with the adaptive focus control from Leica. Images were analysed with ImageJ software. Biotin (1:1 dilution, 80 μM stock solution) was added immediately prior to imaging and images were taken every minute for 1 h. Movies obtained from the time-lapse images were processed using the ImageJ plugin HyperStackReg V5.6. (Ved Sharma) and the fluorescence intensity decay due to photobleaching was corrected using the Exponential Fitting method from the ImageJ Bleach Correction tool (https://f1000research.com/articles/9-1494/v1). The TNFα_GFP mean fluorescence intensity was quantified from images taken at 0, 10, 20, 30, 40, 50 and 60 minutes and plotted using GraphPad Prism 8 software.

### Immunofluorescence and confocal microscopy

2*10^5 day 3 moDCs were seeded on a coverslip and 72 h later treated with LPS (1 μg/ml) alone or in combination with Brefeldin A (10 μg/ml) and Monensin (2 μM), or with Bafilomycin (100 nM) for 6 hours. DMSO was used as negative control. After this time, cells were fixed in either 4 % paraformaldehyde (PFA) or in 100 % methanol and subsequently incubated for 20 minutes in 0.1 M quenching solution (NH4Cl in PBS). Cells transfected with the TNFα plasmid were also seeded on coverslips using Leibovitz’s medium for 3 h prior to being treated with biotin (1:1 dilution) for 30 min. After this time, cells were fixed with 4 % PFA. PFA-fixed cells were then washed twice with PBS and permeabilised in permeabilisation buffer (2.5 % Bovine Serum Albumin, 1 % donkey serum and 0.15 % Triton X-100) for 30 minutes prior to being incubated o/n and at 4 °C with specific antibodies against TNF-α (MACS miltenyi biotec, #130-091-649), Vamp-3 (Abcam #Ab5789, 1:200 final dilution, the ER marker PDI (Novus Biologicals #NBP2-02082), the Golgi cis/trans marker GM130 (BD transduction laboratories #610822), the early endosomal marker EEA1 (BD Biosciences #610456/57, 1:100 final dilution) and galectin-9 (R&D biosystems, #AF2425, 1:40 final dilution). Methanol-fixated cells were washed, permeabilised and incubated o/n at 4 ºC with antibodies against Vamp-3 and galectin-9 (as per previous staining protocol). Please note that methanol-fixed cells show galectin-9 nuclear staining as an artefact of the fixation process, which does not represent real localisation of the protein.

Cells were afterwards washed extensively prior to being incubated for 1 h at room temperature with a donkey-anti-mouse alexa 488 secondary antibody (Thermofisher, Massachusettes, US, A21202, 1:400 final dilution) and a donkey-anti-goat alexa 568 (ThermoFisher Scientific, #A11055, 1:400 final dilution) in permeabilisation buffer. PFA and methanol fixed cells were then washed and nuclei stained with DAPI for 10 minutes at room temperature. Coverslips were embedded in glass slides using Mowiol prior to being stored at 4 °C until imaging. Samples were imaged with both a Leica DMI6000 epi-fluorescence microscope fitted with a 63x 1.4 NA oil immersion objective, a metal halide EL6000 lamp for excitation, a DFC365FX CCD camera and GFP and DsRed filter sets (all from Leica, Wetzlar, Germany) as well as a Zeiss LSM900 confocal laser scanning microscope. For the images acquired with the Leica DMI6000 epi-fluorescence microscope we used a 63x 1.4 NA oil immersion objective, a metal halide EL6000 lamp for excitation, a DFC365FX CCD camera and GFP and DsRed filter sets (all from Leica, Wetzlar, Germany). Focus was kept stable with the adaptive focus control from Leica. For the images acquired with eh Zeiss LSM900 microscope we used a 63x EC Epiplan-NEOFLOUAR oil immersion objective and three laser module URGB (405, 488, 561, 640 nm). Images were analysed with ImageJ software. Fluorescence overlap was quantified using an unbiased macro in ImageJ which automatically selected imaged cell areas based on fluorescence intensity. The Manders correlation coefficients were calculated using the JACoP plugin and the line scan graphs representing the fluorescence cross-sections were quantified using the Plots profile command in ImageJ and depicted using GaphPad Prism 8 software. The cross-sections were established by drawing a perpendicular line from the nuclear membrane 20-250 microns (for Vamp-3 intensity quantification or the colocalization analysis, respectively) towards the plasma membrane furthest away from the nucleus.

Image analysis from experiments using the TNFα RUSH construct was performed using ImageJ. Integrated density of Vamp3 in the PD1, GM130 and EEA1 channels was quantified using an unbiased macro in ImageJ which masked the fluorescence intensity in each subcellular compartment after applying the Yen, Default and Yen thresholding algorithms (for the PD1, GM130 and EEA1 channels respectively). Each mask was saved as a region of interest (ROI), transferred to the Vamp3 channel, and the integrated fluorescence density from each ROI was quantified.

### Statistical analysis

The statistical test used to analyse each data set is described in the corresponding figure legend. Unpaired t-test was used to compare mean values between NT and *gal9* siRNA transfected cells. An anova-two way was used to quantify Vamp-3 intensity across the cytosol.

## Acknowledgements

We thank Franck Perez for depositing the TNFα_EGFP RUSH construct to Addgene. We thank Pascal W. T. C. Jansen for his assistance with the proteomics experiments. We thank Geert van den Bogaart for critical reading of the manuscript and helpful discussion. This work is supported by grant 11618 from the Dutch Cancer Society. C.G Spruijt and Michiel Vermeulen are part of the Oncode Institute, which is partly funded by the Dutch Cancer Society. Annemiek van Spriel is supported by the Netherlands Organization for Scientific Research NWO Gravitation Programme 2013 grant (ICI 000-23), ZonMW (project 09120012010023), the Dutch Cancer Society (KWF) (12949/2020), and was awarded a European Research Council Consolidator Grant (Secret Surface, 724281).

## Figure legends

**Supplementary figure 1. Endogenous levels of galectin-9 in moDCs can be modulated.** Day 3 moDCs were transfected with *gal9* siRNA or a NT siRNA. Levels of galectin-9 were assessed by flow cytometry 48 h after transfection. **A.** Graph shows galectin-9 levels at the plasma membrane. **B.** Histogram depicts total protein levels, both surface-bound and intracellularly. As shown, galectin-9 was depleted both at the plasma membrane as well as intracellularly. NT (black line, unfilled population), *gal9* siRNA-transfected moDCs (light grey population). Black dotted line represents isotype control values. Graphs are representative for one donor and numbers in inset indicate geometric mean fluorescence intensity (gMFI). **C.** Total lysates from NT and *gal9* siRNA transfected cells were subjected to Western Blot and Galectin-9 expression was analysed. Tubulin was used as loading control.

**Supplementary Figure 2. Several stimuli enhance the secretion of specific cytokines. A.** NT siRNA and *gal9* siRNA previously transfected moDCs were seeded on day 6 and treated with either 1 μg/ml LPS, 4 μg/ml R848, 1 μg/ml poly I:C or *Candida albicans* (*C. albicans*, ratio 1:10) for 6 (LPS-stimulated cells) or 16 hours (all other stimuli) prior to supernatants being collected. Specific ELISA reactions for the above cytokines were performed. Data is representative for one donor. **B.** Day 3 moDCs were transfected with NT siRNA or *gal9* siRNA. Forty-eight hours after transfection, cells were treated with LPS, R848 or zymosan particles for the indicated time points and secreted levels of TNFα, IL-6, IL-10 and IL-12 determined by ELISA. Graphs show representative data from one donor.

**Supplementary Figure 3. Cytokines accumulate intracellularly upon galectin-9 depletion. A.** Additional representative images of the results shown in Figure 2C. DAPI, TNFα and galectin-9 immunofluorescence stainings from NT (above) and *gal9* siRNA (below) transfected moDCs treated for 6 hours with 1 μg/ml LPS are shown. Scale bar: 10 μm.

**Supplementary Figure 4. Galectin-9 depleted moDCs eliminate intracellular cytokines via lysosomes**. Additional representative images of the results shown in Figure 3C. DAPI, TNFα and galectin-9 immunofluorescence stainings from NT (above) and *gal9* siRNA (below) transfected moDCs treated for 6 h with 1 μg/ml LPS and Bafilomycin (200 μM) are shown. Scale bar: 10 μm.

**Supplementary Figure S5. Cytokines do not traffic after the Golgi complex in galectin-9 depleted moDCs. A.** Additional representative images of the results shown in Figure 4C. DAPI, TNFα and EEA1 immunofluorescence stainings from NT (above) and *gal9* siRNA (below) transfected moDCs treated for 6 hours with 1 μg/ml LPS are shown. Arrows indicate sites of colocalisation between TNFα and EEA1. **B.** NT and *gal9* siRNA moDCs were treated with 1 μg/ml LPS in combination with the Golgi inhibitors Brefeldin A (BrefA, 10 μg/ml) and Monensin (Mon, 2 μM) after which cells were fixed and stained with specific antibodies against TNFα, the Golgi marker GM130 and DAPI for nuclear staining. Representative airyscan confocal images of three independent experiments are shown. Scale bar: 10 μm. Graphs: fluorescent cross-sections as indicated.

**Supplementary movie 1. Wild type DC TNFα-GFP trafficking dynamics.**

Movie corresponding to Figure 5B

**Supplementary movie 2. Galectin-9 depleted DC TNFα-GFP trafficking dynamics.**

Movie corresponding to Figure 5B

