## Supplementary figures and images for "Galectin-9 interacts with Vamp-3 to regulate cytokine secretion in dendritic cells"

### Supplementary Figure 1

**Supplementary Figure 1**

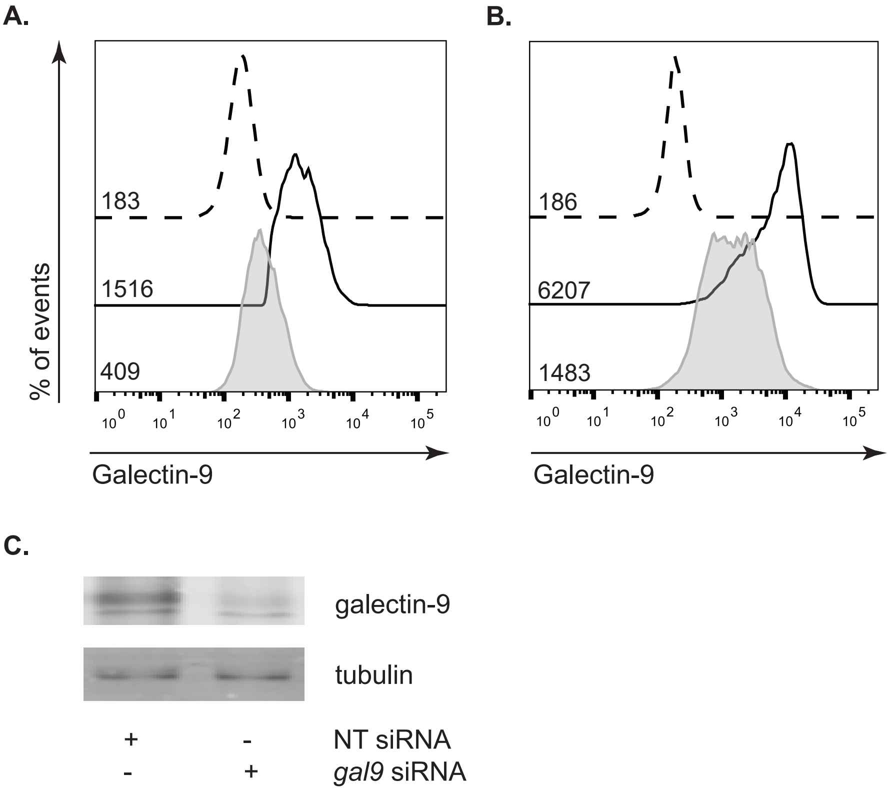

### Supplementary Figure 2

**Supplementary Figure 2**

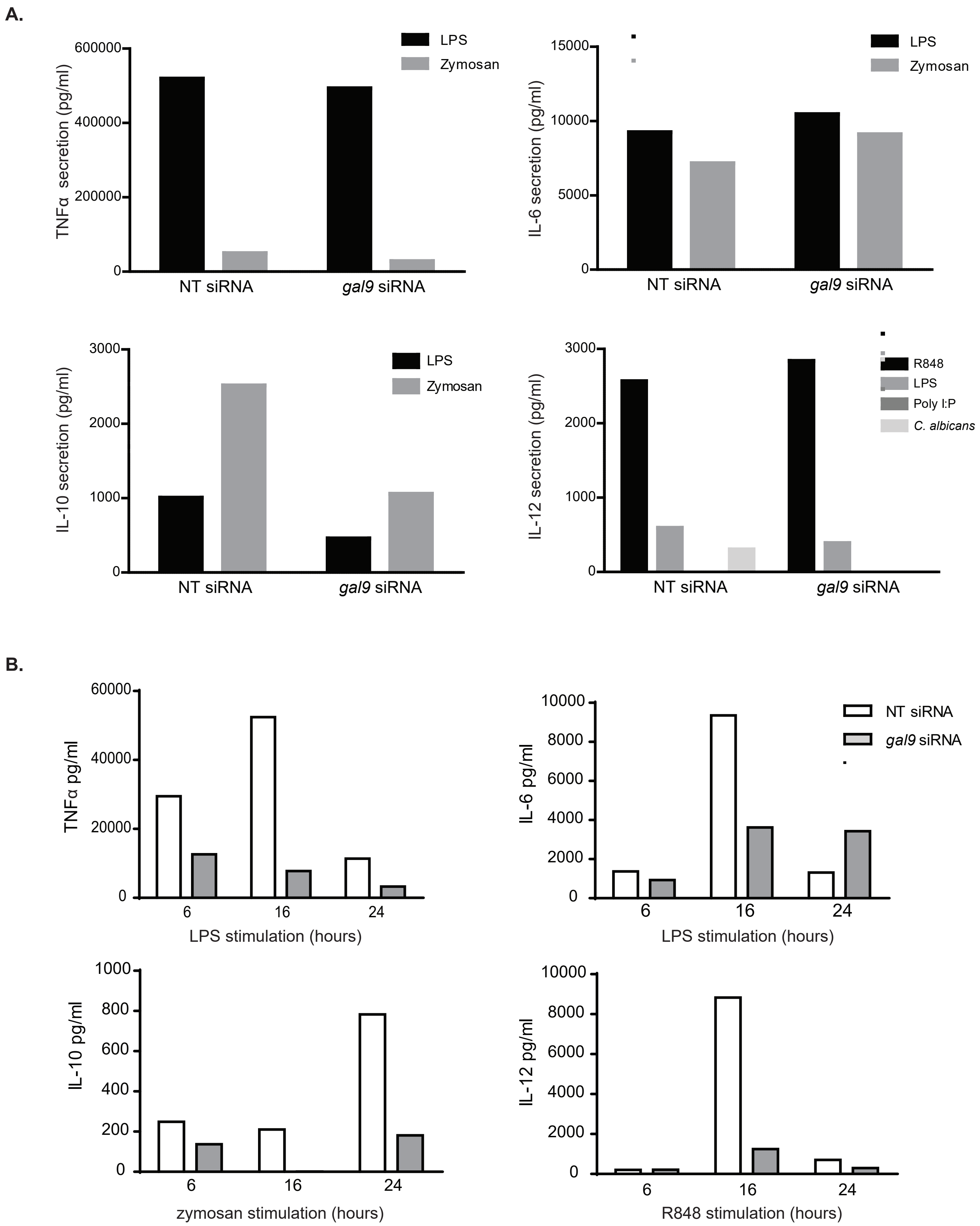

### Supplementary Figure 3

**Supplementary Figure 3.**

**A.**

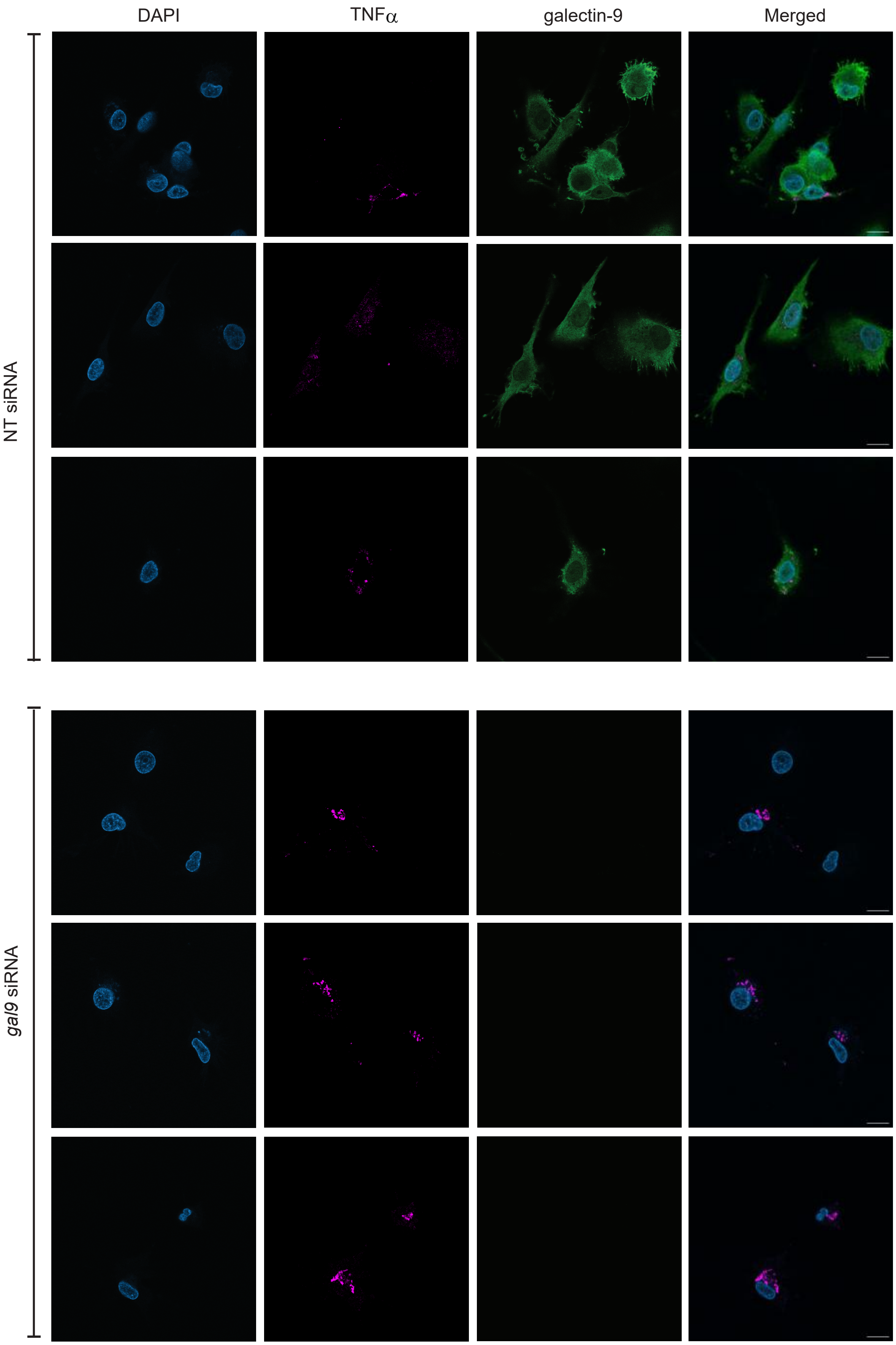

### Supplementary Figure 4

**Supplementary Figure 4**

**A.**

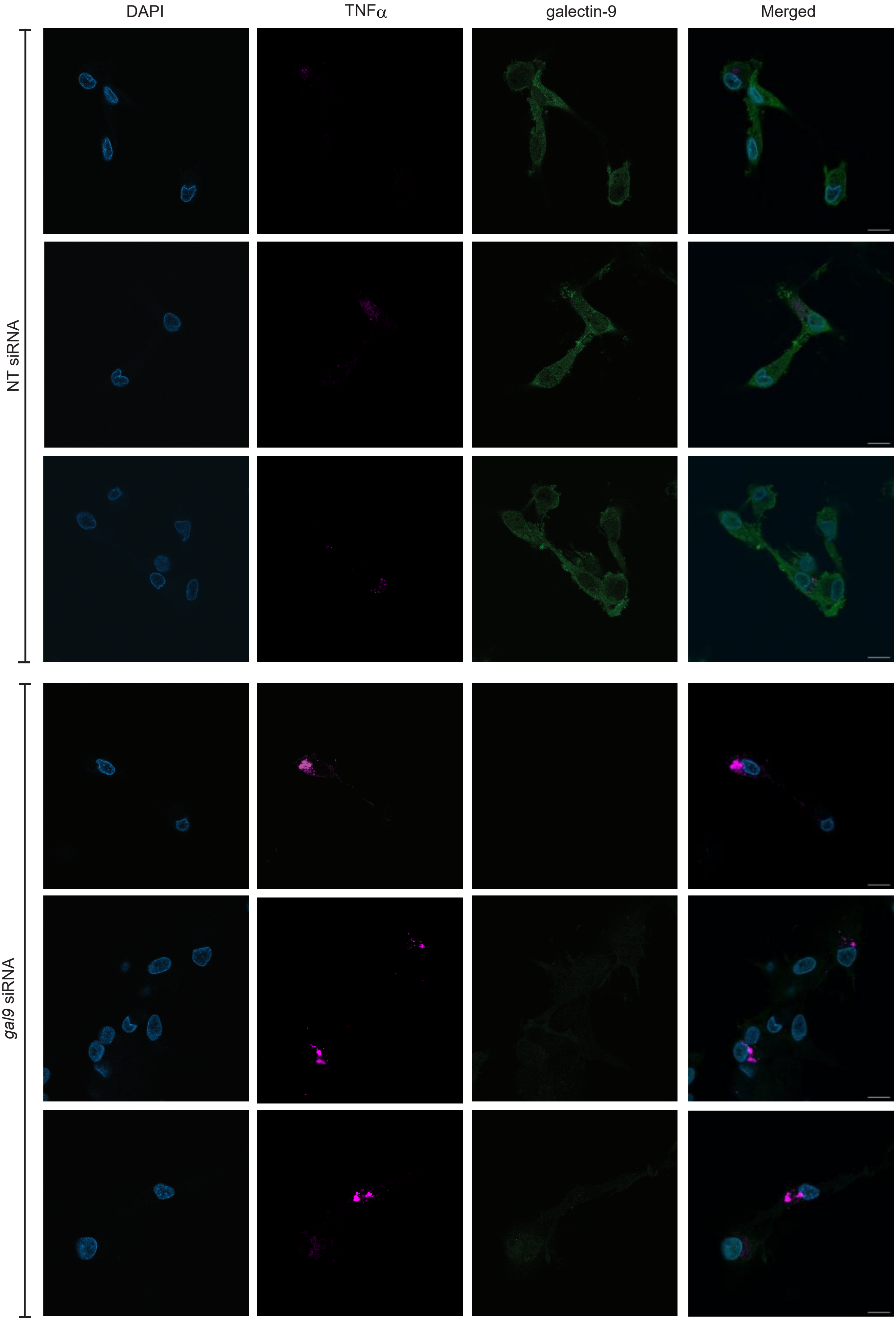

### Supplementary Figure 5

**Supplementary Figure 5.**

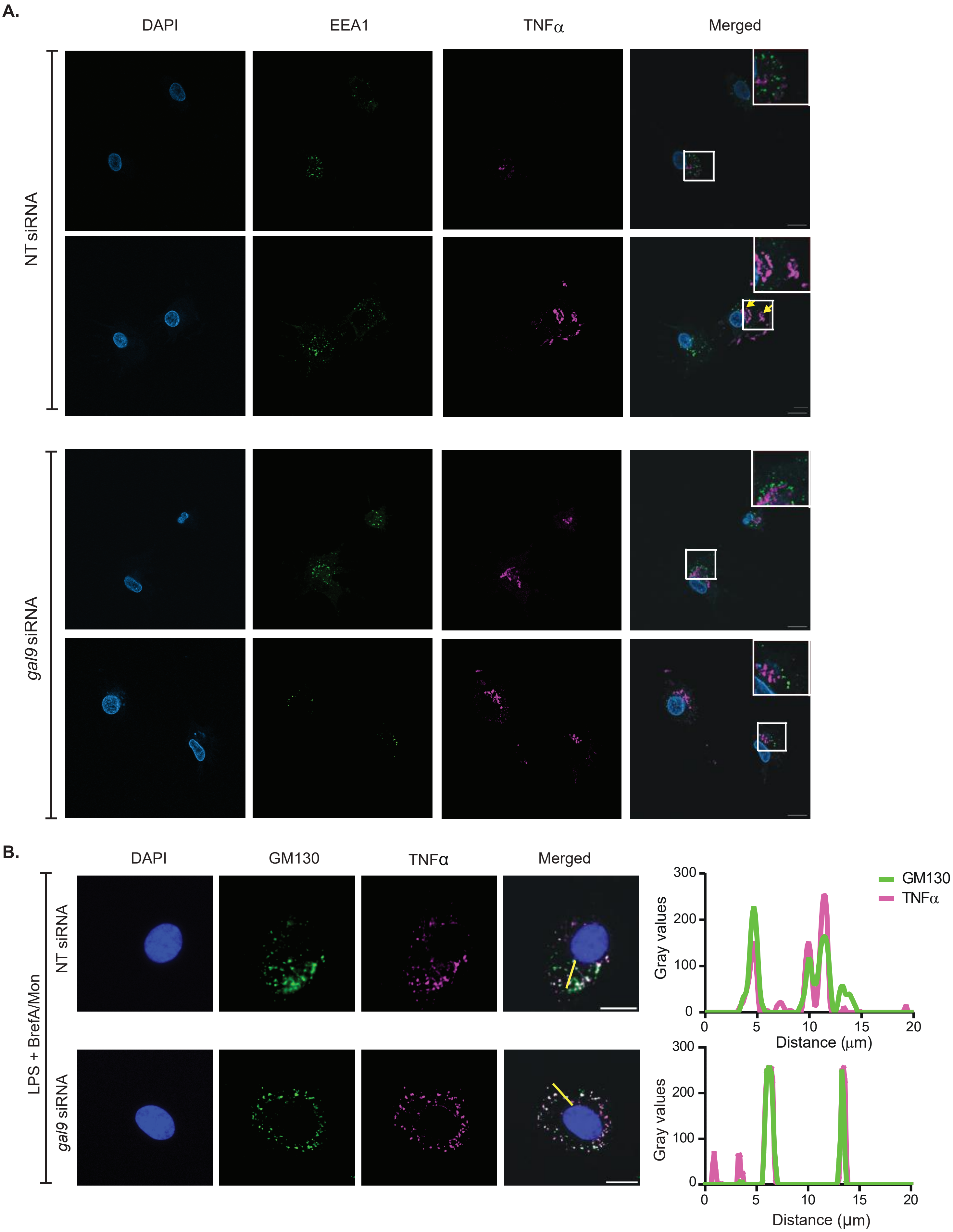
